# A general high-throughput mucus microrheology platform for quantitative and scalable mucus phenotyping

**DOI:** 10.1101/2025.01.09.632077

**Authors:** Feng Ling, Ayşe Tuğçe Şahin, Bernardo Miller Naranjo, Stefano Aime, Doris Roth, Anna Pukaluk, Niels Tepho, Romina Augustin, Benedetta Zampa, Andrea S. Vendrame, Mark-Christian Klassen, Ellen Emken, Marion Kiechle, Mircea Gabriel Stoleriu, Pontus Mertsch, Ruth Olmer, Yohannes Tesfaigzi, Oliver Lieleg, Janna C. Nawroth

**Author notes:** To whom correspondence should be addressed: Janna Nawroth.

## Abstract

Mucus mechanics regulate barrier, clearance, and transport functions across epithelial tissues, yet quantitative rheology remains difficult to scale because available assays require large volumes, specialized equipment, or extensive manual analysis. Here, a high-throughput microrheology workflow based on Differential Dynamic Microscopy is established for frequency-dependent measurements from 3–10 µL mucus samples and intact mucosal surfaces using standard epifluorescence microscopy. The workflow combines preloaded tracer chambers, controlled sample handling, short video acquisition, and automated analysis to reduce operator intervention while preserving sensitivity to mucus relaxation on 1–20 s time scales. Validation against shear rheometry and particle tracking in reconstituted mucin gels shows robust recovery of viscoelastic trends across mucus-like material states. Applications in human airway epithelial air–liquid interface cultures resolve culture-medium, age, donor, COPD, cigarette-smoke, cystic-fibrosis treatment, and IL-13-associated differences, including spatial heterogeneity in situ. Measurements of clinical cervical mucus further demonstrate parallel testing of scarce specimens and sensitivity to osmotic swelling. This platform provides a scalable route to quantitative mucus phenotyping for disease modeling, therapeutic testing, and future precision-medicine studies.

## 1 Introduction

Healthy human mucus is a multifunctional material for epithelial protection. It acts as a barrier to pene-tration;^[1–8^ ^]^ mediates transport;^[9–14^ ^]^ provides lubrication;^[15–17^ ^]^ and preserves epithelial hydration.^[18–20^ ^]^ These functions are crucially dependent on mucus rheology, i.e., how mucus deforms under applied force. While mucus mostly consists of water, its small fraction of solid components (e.g., 5% mucins, 2% lipids, 1% salts, and 0.02% DNA and other molecules in respiratory mucus^[8,21–23^ ^]^) generate complex frequency-and scale-dependent viscoelastic behaviors that regulate epithelial barrier functions by controlling the movement and diffusion of particulate matter,^[9,24–29^ ^]^ as well as enabling dynamic mucus clearance.^[10–12,30^ ^]^ The importance of these behaviors is particularly evident in the respiratory system, where altered mucus properties are associated with higher risks of infections and muco-obstruction, for example in asthma, chronic obstructive pulmonary disease (COPD), cystic fibrosis (CF), primary ciliary dyskinesia (PCD), and related disorders.^[5,31–35^ ^]^ In the human airway epithelium, a mucus layer secreted by specialized glands and secretory cells interspersed between multiciliated cells forms the mobile substrate for the trapping and removal of pathogens and debris.^[7,30,36^ ^]^ Fundamentally, effective clearance of trapped substances relies on the interplay between cilia activity and mucus rheology.^[8,37^ ^]^ For instance, reduced mucus viscosity disrupts ciliary coordination and impairs globally directed flow.^[10^ ^]^ On the other hand, mucus with high viscoelasticity can also hinder ciliary motion, resulting in reduced mucociliary clearance and mucus plugging.^[38,39^ ^]^ Because of this complexity, the biophysical mechanisms and physiological implications of changes to mucus rheology remain poorly understood, limiting our understanding of how abnormal mucus rheology contributes to the onset and progression of chronic airway diseases.^[5,37^ ^]^

Thus, it is essential to develop tools that can examine mucus viscoelastic properties and track their evolution over the course of disease onset, progression and treatment under physiologically relevant conditions. Recent advances have enabled patient-specific disease modeling using human airway epithelial cultures and Organ-Chip models at air-liquid interface (ALI).^[40–44^ ^]^ Such models have helped reveal that abnormal numbers and phenotypes of mucus-secreting cells are key indicators of many chronic airway diseases, including asthma, COPD and bronchiectasis.^[7,45^ ^]^ Further, certain genetic and epigenetic variations, pre-existing conditions and environmental stressors may alter the phenotype or the abundance of mucus-secreting cells and raise the risk for developing chronic lung disease, such as COPD.^[46–48^ ^]^ However, beyond correlations, missing thus far is a mechanistic understanding of how changes in mucus secretion and rheology drive disease progression, and why some individuals worsen after pollutant exposure while others remain resilient.^[49–53^ ^]^ Moreover, therapeutic agents intended to penetrate or restore the mucus barrier require quantitative assays that characterize mucus viscoelastic and structural properties.^[54,55^ ^]^ Similar challenges extend to mucosal tissues in digestive and reproductive organs, where the mechanical properties of mucus are essential for barrier, clearance, and lubrication functions.^[14,56,57^ ^]^ In all these systems, research has been constrained by the lack of high-throughput methods capable of analyzing large numbers of small volumes (microliters) of mucus provided by experimental models, such as in vitro cultures, Organ-on-chips systems,^[58^ ^]^ animal models, as well as clinical biopsies.

While the clinical significance of mucus and sputum rheology has been recognized since the late 1960s, early measurement devices using magnetically actuated spheres with ∼ 100 µm radius offered only modest precision and narrow frequency range.^[9,26,49,59–61^ ^]^ These limitations motivated the adoption of Multi-Particle Tracking (MPT) microrheology, which infers mucus microstructure and viscoelasticity from broad-spectrum thermal fluctuations of micrometer-sized particles.^[8,62–65^ ^]^ Although microfluidic implementations of MPT (µ^2^-rheology) have enabled high-throughput screening of synthetic hydrogels pre-mixed with tracer beads,^[66^ ^]^ direct application of this method to mucus samples has not been achieved, likely due to constraints in handling of small mucus volumes. Moreover, MPT typically involves multiple manual steps that are difficult to implement in high-throughput settings.^[67^ ^]^ For example, imaging contrast needs to be carefully optimized at acquisition and analysis to reconstruct accurate and precise particle trajectories. Manual intervention is also frequently necessary during analysis to filter out spurious trajectories based on intensity, track length, and location. Such manual workflows make it nearly impossible to monitor in vitro rheological dynamics at time resolutions relevant to secretory cell responses and mucus composition changes in response to pollutants or drugs, which often occur within less than 48 hours. Likewise, scaling MPT for industrial and large-cohort studies, where hundreds of samples must be analyzed reproducibly and free of user bias, remains impractical.^[68^ ^]^ Together, these limitations have prevented the systematic, large-scale quantification of mucus rheology across conditions, donors, and time—leaving a critical gap between experimental models and clinically actionable measurements. Differential Dynamic Microscopy (DDM) offers a promising alternative.^[69^ ^]^ Like MPT, DDM can measure viscoelastic moduli across a wide range of frequencies using only a light microscope, a digital camera, and conventional illumination methods. In stark contrast to MPT, however, DDM analysis can be fully automated without any user input, making it far more suitable for high-throughput experiments that demand automation and standardization.^[68,70–72^ ^]^ DDM also offers enhanced statistical precision and allows for the study of particles smaller than the optical resolution limit since it does not rely on tracking specific features across image stacks. Combined with its ability to work with low-signal and optically dense materials,^[72–76^ ^]^ DDM is particularly suitable for large-scale studies of micro-liter *in vitro* mucus samples containing variable mucin concentrations and cellular debris.

Although DDM can directly measure particle diffusion rates in mucus from bright-field videos alone,^[77^ ^]^ the presence of cell debris and particulate matter with unknown size distributions (*e.g.*, residuals from cigarette smoke exposure) makes it difficult to standardize analysis across different experiments. Therefore it is still advantageous to work with mono-dispersed fluorescent tracer beads of known size, despite the risk of altering the original viscoelastic characteristics of micro-scale mucus samples due to mixing and dilution. The long viscoelastic relaxation time of mucus also implies that video footage over multiple minutes needs to be acquired per region of interest according to conventional DDM practices.^[22,73,78,79^ ^]^ Data handling challenges aside (at 100 frames per second, a minute-long 16-bit uncompressed video generates almost one gigabyte of data even at 256 × 256 pixels), these long acquisition windows inherently risk motion artifacts from microscopic flow, drift, and vibrations in common microscopy set-ups.^[80^ ^]^

In this study, we overcome these challenges by advancing and integrating state-of-the-art DDM methodologies together with optimized sample collection and preparation steps. In multiple proof-of-concept applications ranging from *in vitro* respiratory cultures to clinical cervical samples, we demonstrate the power of fluorescence-based DDM for the high-throughput analysis of small (3–10 µL) mucus droplets as well as for in situ analysis of mucosal surface where the mucus secretions is too sparse for collection. Our technology can be employed by non-experts with access to standard epifluorescence microscopy. We highlight practical considerations and specific pain points for deployment and discuss how our platform and its derivatives can benefit future translational research.

## 2 Results

### 2.1 A Scalable High-Throughput Microrheology Platform

#### Throughput capacity and human attention costs

Our platform (**Figure 1A**) can analyze as little as 3 microliter mucus samples using only a few minutes of sustained human attention per sample, not counting the time required for laboratory preparations and automated analysis done per experimental run. Two preparation steps precede an experiment: (i) fabrication of capillary chambers pre-loaded with evaporation-dried fluorescent tracers, and (ii) calibration of tracer hydrodynamic radius to correct for changes introduced by the evaporation-rehydration cycles (see Experimental Section and Supporting Information). For DDM acquisition, we load one mucus sample per chamber and gently re-suspend the surface-bound tracers, and record epifluorescence videos from 3-10 regions of interest (ROIs) based on observed heterogeneity of particle motion (typically 5 ROIs spanning the chamber are used). If coexisting phases are evident (*e.g.*, liquid pockets inside denser gel condensates), each phase is analyzed separately for consistency. To limit stage or flow artifacts while maintaining throughput, we typically acquired 20-second videos at 100 frames per second (fps). 500 fps were sometimes used for lower-viscosity mucus samples that require greater temporal resolution, and up to 1000 fps were used for hydrodynamic radius calibration experiments; see Supporting Information for more details and the full optical setup. This predefined time-frequency window is a deliberate trade-off between statistical power and required experimental effort, as extending measurements to very high frequencies or long videos increases data-acquisition, storage and processing demands without necessarily yielding additional rheological information for DDM analysis. With DDM run unattended in batch after imaging, the only throughput-limiting steps are sample collection, loading and image acquisition.

**Figure 1:**
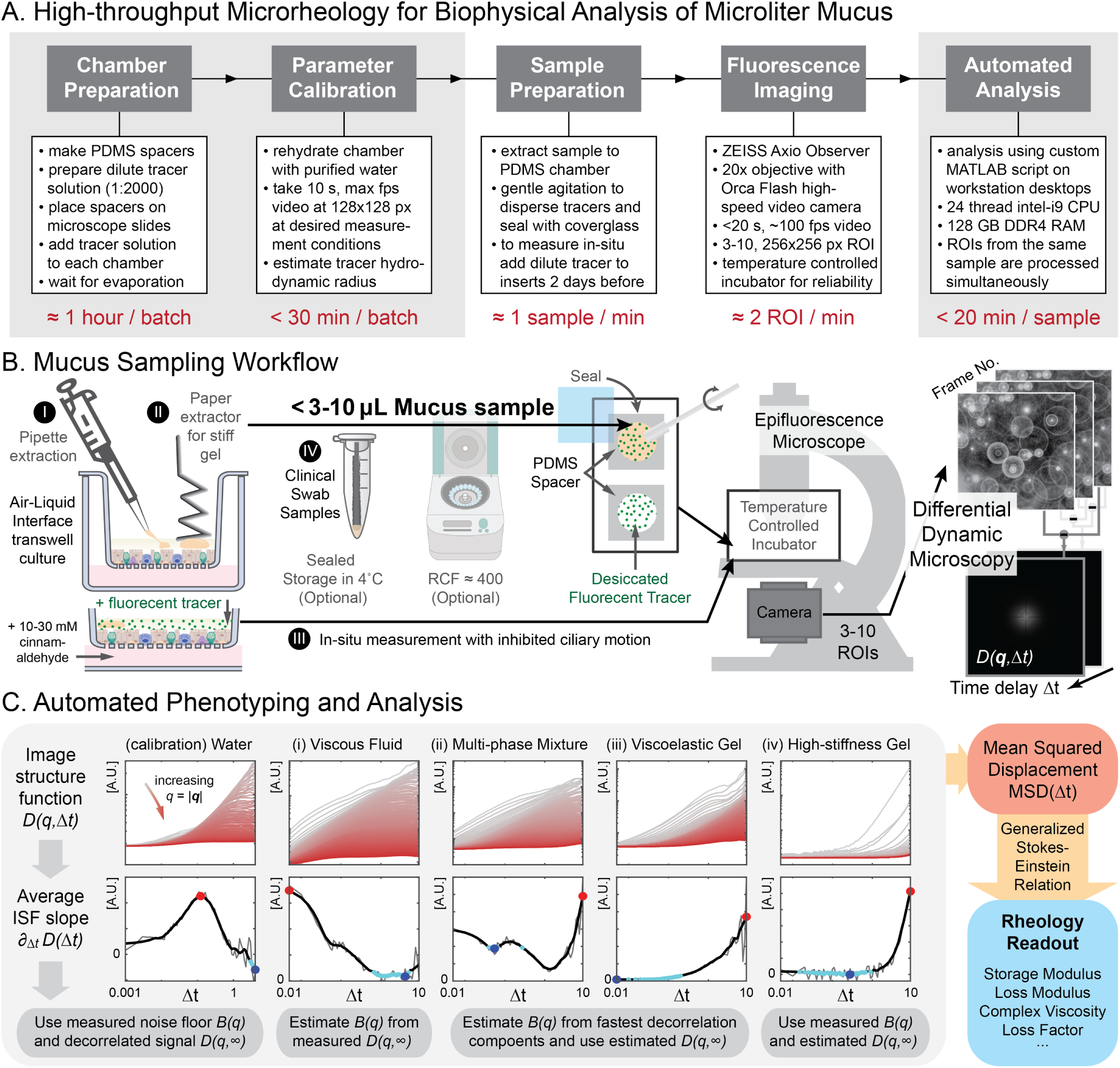
High-throughput mucus microrheology workflow. **A.** Our workflow takes approximately 30 min to analyze every 3-10 µL mucus sample, requiring only a few minutes of sustained human attention outside of laboratory preparation work. In a trial run, 60 samples were prepared in 62 min, and 235 regions of interest were imaged in 123 min and analyzed unsupervised before the next working day. **B.** Depending on the mucus state and secretion source, mucus viscoelasticity was measured either from apical transwell extractions (I) or (II), in-situ with ciliostasis induced by cinnamaldehyde-infused basal medium (III), or clinical sources (IV). The extracted droplets can be sealed in parafilm-wrapped Eppendorf tubes if transportation or storage is necessary before analysis. Low speed centrifugation can help separate mucus droplets from buffer solutions. Measurement chambers are made from Polidimethylsiloxane (PDMS) spacers with small round holes sandwiched between glass slides and cover glass. Before loading a sample, dilute fluorescent tracer solution is filled to each chamber and let dry in a dark box until solvent evaporation. After verifying bead integrity and hydrodynamic radius via rehydration tests, mucus droplets are transferred per chamber, gently mixed and sealed for epifluorescence microscopy inside a temperature-controlled incubator for Differential Dynamic Microscopy (DDM) analysis. **C.** Test samples are automatically typed into different viscoelastic categories (i–iv) based on qualitative features of the image structure function (ISF) and its time derivative. Noise floor and decorrelation signal of the ISF can then be estimated to obtain mean squared displacement (MSD) and to reconstruct desired rheological readouts in sequence. Transwell and centrifuge illustration created with BioRender https://BioRender.com/d80d467.

To test our throughput capacity, we conducted a trial run with 60 individual mucus extractions. Preparation of capillary chambers and desiccated tracers took less than 10 minutes of labor with about 1 hour of waiting, while sample loading and mixing required ∼ 1 min per sample (62 min total). Imaging steps (locating tracers and focusing) took ∼ 1 min for 2 ROI (123 min for 235 ROI). The subsequent DDM analysis took ≤ 20 min per sample on a desktop computer overnight in batch mode without human supervision. In total, the aggregated human attention cost for this 60-sample trial is less than 4 hours, and the rest of the analysis routine (< 20 h) is finished by the next working day. Compared with a viscosity-only commercial microfluidic screen (96-sample in 24 hours), our approach, even as implemented in academic laboratory conditions, achieved comparable throughput with minimal human attention costs.^[81^ ^]^

### Sample collection and in-situ imaging

Given mucus variability across tissue type, donor, culture protocol, and disease state, as well as the susceptibility of mucus viscoelasticity to perturbations such as dehydration, excessive shear and pH, we standardized multiple sample collection and imaging strategies tailored to different experimental conditions and objectives. Specifically, we measured mucus collected from ALI cultures via conventional pipetting or custom paper-based extractors, or from human donors via clinical swabs in custom capillary chambers. We also performed direct in-situ measurements in live ALI culture with inhibited ciliary motion to gain additional insights on the spatial distribution of mucus and enable measurements when mucus was too sparse for collection. All strategies are outlined in the following; for details see Materials and Methods. (I) Conventional pipette extraction (Figure 1B(I) and Video S1, Supporting Information). To avoid accidental collection of buffer or culture medium, we first verified the elastic stretching response of each droplet before extraction. Pipette volume was set to a smaller value than the droplet size such that droplets remained attached to the tip opening rather than being fully engulfed. This also minimized wasteful sample adhesion to the pipette tip wall during transfer. For cases where the mucus droplets were strongly tethered to the cell surface, a small volume of PBS (∼27 µL/cm^2^) was added to *all* test conditions 3 min before extraction to loosen adhesion without significantly altering the relative viscoelastic differences. If buffer was collected or mucus droplet became trapped inside the tip, mild centrifugation at 400 relative centrifugal force (RCF) for 10 min at 4^◦^C was used to separate mucus from the supernatant before transferring sample into the capillary chamber. (II) Paper-assisted extraction (Figure 1B(II) and Video S1, Supporting Information). In experiments where mucus droplets were highly viscous and measurement without any buffer dilution was desired, UV-disinfected weighing paper was cut and folded into strips to lift off stiff mucus droplets instead. This approach was able to lift off accumulated mucus while preserving cell layer integrity better than repeated pipette collection, which required much finer operator control. (III) In-situ imaging (Figure 1B(III)). To probe spatial heterogeneity of mucus droplets, or when secreted mucus layer was too thin to be collected manually (no accumulated semi-transparent mucus bump visible in the insert), we performed direct in-situ imaging on the live cell surface after transient inhibition of ciliary beating. While this approach avoids mucus extraction and loading, it requires introducing fluorescent tracers in buffer to the apical side of ALI culture. To mitigate any dilution artifacts, the buffer volume was minimized and the tracer suspension was applied at least 24 hours prior to imaging in all measured conditions. (IV) Clinical samples. Patient-derived specimens were collected in the clinic and loaded into the standardized holder in the same manner as in case (I) and (II). Combined with sample loading optimizations discussed above, this approach allowed aliquot-based parallel testing of limited material.

#### Automated sample phenotyping

DDM relies on accurate estimates of the noise floor *B*(*q*) and the decorrelated image structure function (ISF) plateau at near infinite lag *D*(*q,* Δ*t* → ∞) to extract the intermediate scattering function *f* (*q,* Δ*t*) and the corresponding viscoelastic moduli; see Supporting Information for theoretical background.^[72^ ^]^ Because we confined acquisition deliberately to a throughput-friendly window, the identifiability of *B*(*q*) versus *D*(*q,* ∞) becomes predictably regime-dependent: ISF of fast-relaxing, low-viscosity samples densely sample the plateau and favor direct estimation of *D*(*q,* ∞) while restricted frame rates can hinder model-free *B*(*q*) fits; slow-relaxing, stiffer gels can yield robust *B*(*q*) yet might never reach *D*(*q,* ∞) within the duration of the video, requiring informed extrapolations. To minimize human bias and manual effort while remaining robust across orders-of-magnitude viscosity changes, we implement an automated classifier that assigns each sample to one of four categories based on features of the ISF: (i) *viscous-liquid*, (ii) *multi-phase mixture*, (iii) *viscoelastic gel* and (iv) *high-stiffness gel* ; see Figure 1C.

#### Classification logic

We first compute prominent extrema in the *q*-averaged time derivative of the ISF *∂*_Δ_*_t_D*(Δ*t*), then locate their characteristic times, check their ordering, and compare them to the acquisition bounds [Δ*t*_min_, Δ*t*_max_]. (i) If the global maximum occurs at Δ*t* ≈ Δ*t*_min_ and precedes the local minimum, the ISF has reached its long-lag plateau within the window; we classify it as *viscous liquid*. Here *D*(*q,* ∞) is measured directly and *B*(*q*) is fitted under Brownian assumptions and may be frame-rate-limited. (ii) If the first local maximum precedes the global minimum, yet the global maximum sits near Δ*t*_max_, decorrelation of a low-viscosity phase is captured while a more viscous phase remains partially

### 2.2 *DDM Outperforms MPT in High-Viscosity Samples*

correlated at Δ*t*_max_; we classify it as *multi-phase mixture*. (iii) If the first local minimum lies near Δ*t*_min_, and the ISF slope steadily increases without saturation within the window (*D*(*q,* ∞) not observed); we classify it as *viscoelastic gel*, with *D*(*q,* ∞) estimated from periodic pixel-shift or average power spectrum proxies and *B*(*q*) fitted with a fractional Brownian model.^[82^ ^]^ (iv) In the limiting case where the ISF slope remains near zero on average over most of the window and the global maximum occurs at Δ*t*_max_ (*i.e.*, a largely stagnant ISF until the largest lags), we classify it as *high-stiffness gel* ; here *B*(*q*) is read off near the ISF minimum and *D*(*q,* ∞) is again estimated from the shifted or average power spectrum proxies. The fitting procedure and uncertainty propagation of *B*(*q*)-estimation are detailed in Supporting Information Sections 2 and Figure S1.

### 2.2 DDM Outperforms MPT in High-Viscosity Samples

To assess the applicability and reliability of our approach for mucus studies, we first measured the rheology of mucus gels reconstituted from lab-purified porcine gastric mucin (MUC5AC). Lab-purified mucin reconstitutes tend to be better model systems for native mucus than commercial mucin sources;^[83–85^ ^]^ see Materials and Methods. Mucin concentration was chosen to vary from 1 to 4% (w/v) to mimic the range of mucus viscoelasticity we expect to collect in airway cell culture models.^[2,21–23^ ^]^ Samples were processed in pH 4 buffers to maximize the likelihood of viscoelastic gel formation.^[86^ ^]^ **Figure 2** compares results obtained from conventional macrorheology, MPT, and our DDM method.

**Figure 2:**
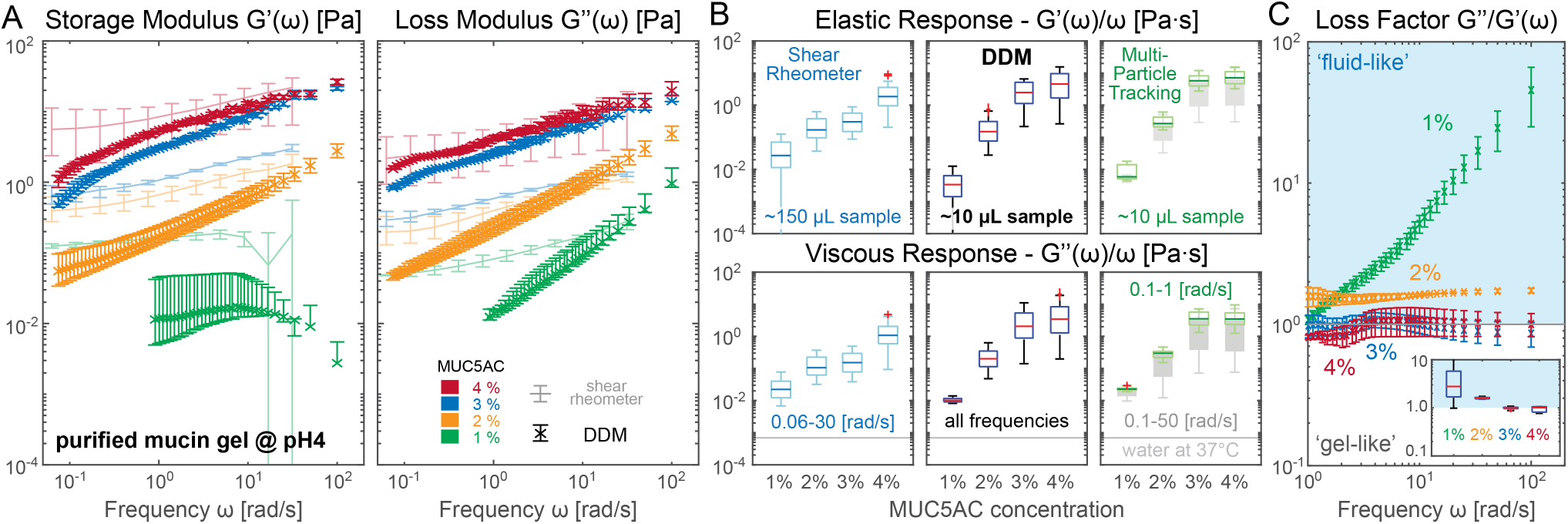
Viscoelastic properties of reconstituted MUC5AC gels. Porcine gastric mucin (MUC5AC) was reconstituted in acidic buffer (pH 4) to simulate mucus gels with physiologically-relevant viscoelastic character. **A.** Storage and loss moduli of 1–4% w/v MUC5AC gels measured using high-throughput DDM (dark cross markers) and conventional shear rheometry (light continuous lines). DDM markers show the median modulus. At each frequency, the lower and upper error-bar limits are the minimum 10th percentile and maximum 90th percentile, respectively, among the *q*-wise estimates from all recorded ROIs. Shear-rheometry error bars show the minimum and maximum of three repeated measurements. **B.** icrorheology output from DDM (middle panels) and Multi-Particle Tracking (MPT, via FIJI Trackmate, right panels) can deviate slightly from shear rheometer results (left panels). DDM microrheology at all frequency ranges closely matches MPT results at restricted frequency (0.1–1 rad*·*s*^−^*^1^, green). MPT results at full frequency ranges (0.1–50 rad*·*s*^−^*^1^, dark gray) produce significantly higher variance especially for high viscosity samples. The light gray line indicates the dynamic viscosity of pure water at measurement temperature. **C.** Average loss factor across different technical replicates with error bars indicating standard error of mean. Samples with higher mucin concentrations behave more “gel-like” (loss factor <1) than “fluid-like” (loss factor >1) near the typical ciliary-beat-frequency range (5–15 Hz). Box plot shows aggregated loss factor for *ω >* 0.5 rad*·*s*^−^*^1^. All measurements were performed in a temperature-controlled sample holder for additional thermal stability.

Figure 2A shows the expected increase of the viscoelastic moduli as a function of mucin concentrations using our high-throughput DDM platform, at a range of frequency responses including typical ciliary beat frequencies of human airway epithelia (5–15 Hz). Corresponding macroscopic shear rheology results are shown as translucent traces behind. To simplify the ranking of the frequency-dependent viscoelasticity of different mucus samples, we discuss the normalized moduli, *i.e.* storage *G*^′^(*ω*) and loss modulus *G*^′′^(*ω*) divided by measurement frequency *ω*, the absolute viscosity |*η*^∗^| = |*G*^′^ + *iG*^′′^|*/ω*, and the loss factor *G*^′′^*/G*^′^(*ω*) as the primary rheological markers. The normalized moduli and absolute viscosity represent the material response in an approximately frequency-independent fashion; see Figure 2B. The loss factor, as the ratio between loss and storage modulus, measures the relative importance of viscous to elastic responses and thus can be used to judge if the sample is more viscous ‘fluid-like’ or elastic ‘gel-like’; see Figure 2C. Importantly, the hydrodynamic radius of tracer particles that scales both moduli is canceled in this ratio, making the loss factor less sensitive to variations in the size uniformity of the tracers.

Both bulk shear rheometry (Figure 2B, left) and DDM microrheology (middle) report an increase in viscoelastic moduli with MUC5AC concentration, but the values do not coincide. Except for the 3% mucin gel, where the smaller 10 µL microrheology sample gave noticeably higher moduli, the shear rheometer consistently reported higher storage moduli than microrheology, whereas the loss moduli remained of similar magnitude. This offset is consistent with a length-scale dependent response of a heterogeneous mucin network. Bulk shear rheometry probes a sample-spanning, load-bearing scaffold and reads a more elastic, gel-like response. In contrast, the microrheology tracers predominantly sample more fluid-like interstitial pores partially coupled to the scaffold. The 3% result, on the other hand, likely reflects sampling of denser or more strongly cross-linked domain rather than a global discrepancy. Together, these observations illustrate that complex biological materials such as mucin gels (or native mucus) can exhibit different mechanical responses when measured as bulk averages (macrorheometer) versus local tracer-based probes that resolve microscopic heterogeneity (microrheology).^[22,37,86–91^ ^]^ This mismatch could also reflect a limitation of one-point microrheology in general; two-point correlation based microrheology methods can in theory reduce sensitivity to probes’ immediate surroundings and general rheological information closer to the bulk measurements.^[86,92^ ^]^

To further validate our platform independent of heterogeneity-induced variabilities, we performed MPT microrheology on the same datasets. Since we chose tracers that can be individually resolved from the same DDM videos, MPT analysis conducted on the same image sequences should produce matching rheology results. Indeed, the MPT results (Figure 2B, right), especially when analyzed at the lower angular frequencies (0.1–1 rad s^−1^, green box plots) and for lower mucin concentration samples (1–2% mucin), closely match our DDM values (middle panels). However, if data from the full angular-frequency range is used (0.1–50 rad s^−1^), MPT generates much larger variances (gray box plots behind green box plots). This discrepancy is due to a reduced signal-to-noise ratio at high mucin concentration where the thermal fluctuation of tracers blends into the high-frequency, static imaging noise that degrades object localization accuracy; see Figure S4 (Supporting Information) for similar test results based on pure synthetic materials. This comparison demonstrates that in a high-throughput workflow with minimal user intervention, DDM is more reliable under suboptimal contrast and signal conditions.

### 2.3 High-Throughput Workflow Enables Longitudinal Studies of Mucus Rheology

Since secretory cell type abundance and composition of *in vitro* ALI human airway epithelial cultures strongly depend on culture medium, donor source and differentiation time points,^[30,42,93^ ^]^ it is expected that the viscoelastic properties of mucus secretions should also vary accordingly. Here, we use this dependence to demonstrate that our platform can track longitudinal changes in mucus rheology across culturing conditions and resolve donor-specific variability in disease-donors such as those derived from COPD patients.

We first compared the viscoelasticities of mucus secreted by primary human airway epithelial cells (pHAEC) cultured at ALI in either PneumaCult ALI medium (PC) or bronchial epithelial growth medium based solution (BEGM-based). **Figure 3A** shows the storage and loss moduli of mucus extracted at multiple time points of differentiation. Our results overlap the previously reported ranges for mucus extracted from PC cultures measured by optical tweezer techniques (Figure 3A, pink bars).^[91,94^ ^]^ Consistent with prior optical tweezer measurements, we also observed a drop in both viscoelastic moduli from samples cultured in BEGM-based media.^[94^ ^]^

**Figure 3:**
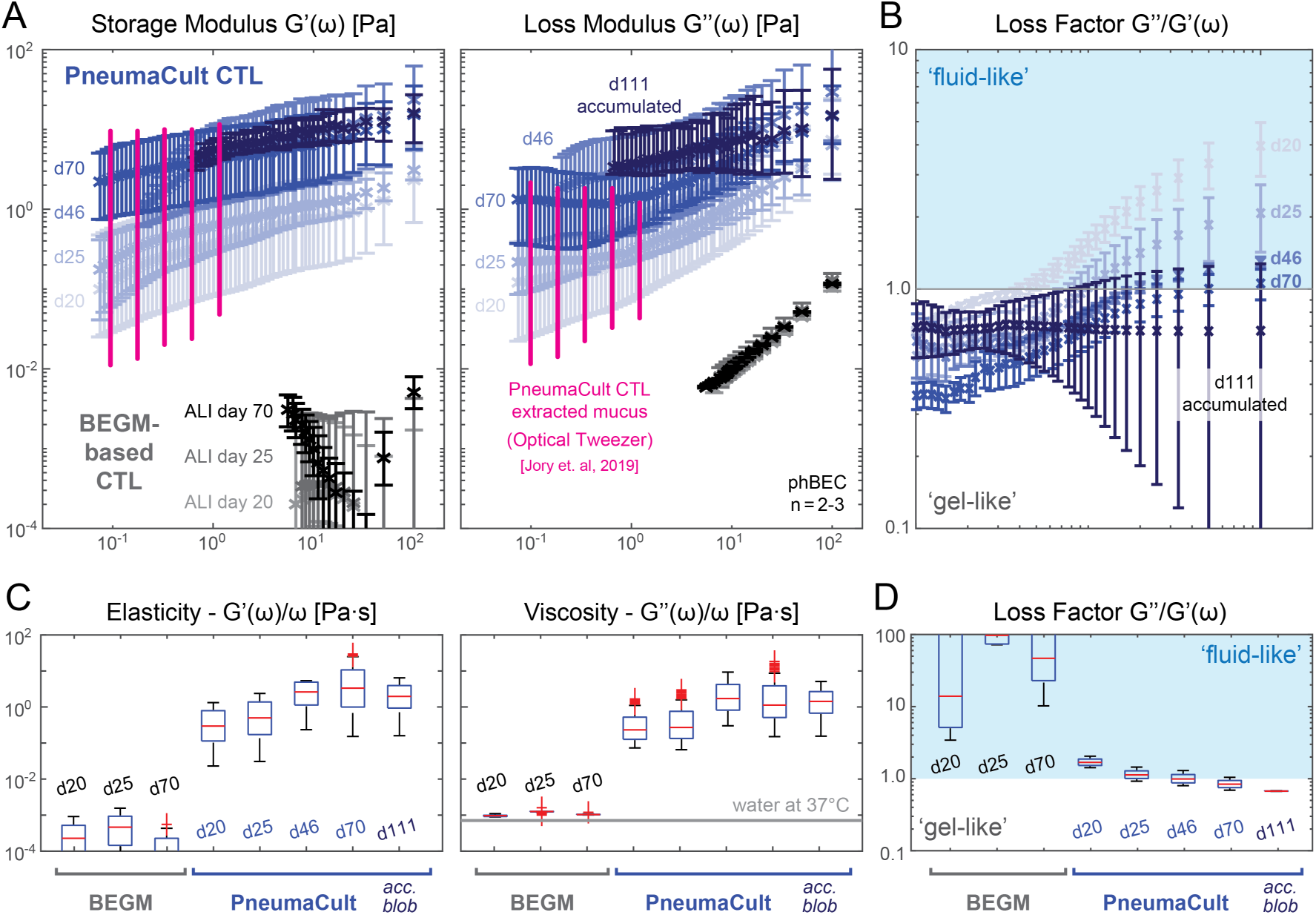
Impact of differentiation medium and culture age on ALI culture mucus viscoelasticity. A–B. Storage and loss moduli (A) and loss factors (B) of mucus extracted at different time points from control pHAEC air–liquid-interface (ALI) cultures maintained in PneumaCult (PC, blue-tinted markers) or BEGM-based medium (grayscale markers). Darker markers denote greater culture age. Pink markers reproduce optical-tweezer measurements of mucus extracted from PC ALI cultures.^[94]^ Except for samples labelled as accumulated (acc. blob), all cultures were washed routinely during maintenance and humidified two days before measurement. Panel A markers show the median moduli. At each frequency, the lower and upper error-bar limits are the minimum 10th percentile and maximum 90th percentile, respectively, among the *q*-wise estimates from all recorded ROIs across the stated technical replicates. Panel B markers show the mean loss factor across technical replicates; error bars indicate the standard error of the mean. **C–D.** Frequency-normalized viscoelastic moduli show that the viscoelasticity of PC-culture mucus plateaued with culture age. BEGM-based mucus consistently showed low viscosities akin to that of water / culture medium. Samples were extracted from *n* = 2–3 same-donor inserts per condition and measured in 2–6 mm-diameter PDMS capillary chambers, depending on the amount secreted. The day-111 accumulated mucus sample was collected using the paper-based extraction device.

Because our platform enables higher-throughput, repeated measurements from small-volume extractions, we monitored how mucus rheology changed with culture age. In PC cultures, mucus viscoelastic moduli increased from day 20 to day 46 at ALI and remained of comparable magnitude through day 70 at ALI (Figure 3C), suggesting mucus properties saturated after epithelial maturation. In contrast, BEGM-

based culture extracts remained weakly viscoelastic (Figure 3A, gray markers), with apparent viscosities close to that of water or culture medium at 37^◦^C (Figure 3C), suggesting that only small amounts of mucin were secreted and/or recovered in the extractions. These results are also consistent with our study showing that PC-based cultures contain a higher proportion of mucus-producing cells than BEGM-based cultures.^[30^ ^]^

In Figure 3B and D, we found that mucus loss factor decreased with culture age and stabilized near or below unity over much of the measured frequency range, indicating a shift toward more gel-like behavior. Note that the substantially more gel-like day-111 PC sample was collected under a different maintenance protocol, where mucus was allowed to accumulate without any routine washing or apical humidification until extraction and measurement. Its significantly lower loss factor is therefore a strong indicator for the importance of implementing consistent control on washing schedule and humidifying dilutions during longitudinal rheological studies.

We next applied the same workflow to PC-based phBEC ALI cultures from three COPD donors with severe disease (GOLD stage IV, groups D or E) and compared them with control PC cultures of similar ALI age (**Figure 4**). Visible ciliary beating appeared later in the COPD donor cultures than in controls (∼ 28–35 days after air lift compared with ∼ 15 days), consistent with reported differentiation defects;^[95^ ^]^ however, mucus rheology did not shift uniformly across COPD donors. Donor #1 produced high-modulus mucus with loss factors near or above unity, donor #2 overlapped the control loss-factor

**Figure 4:**
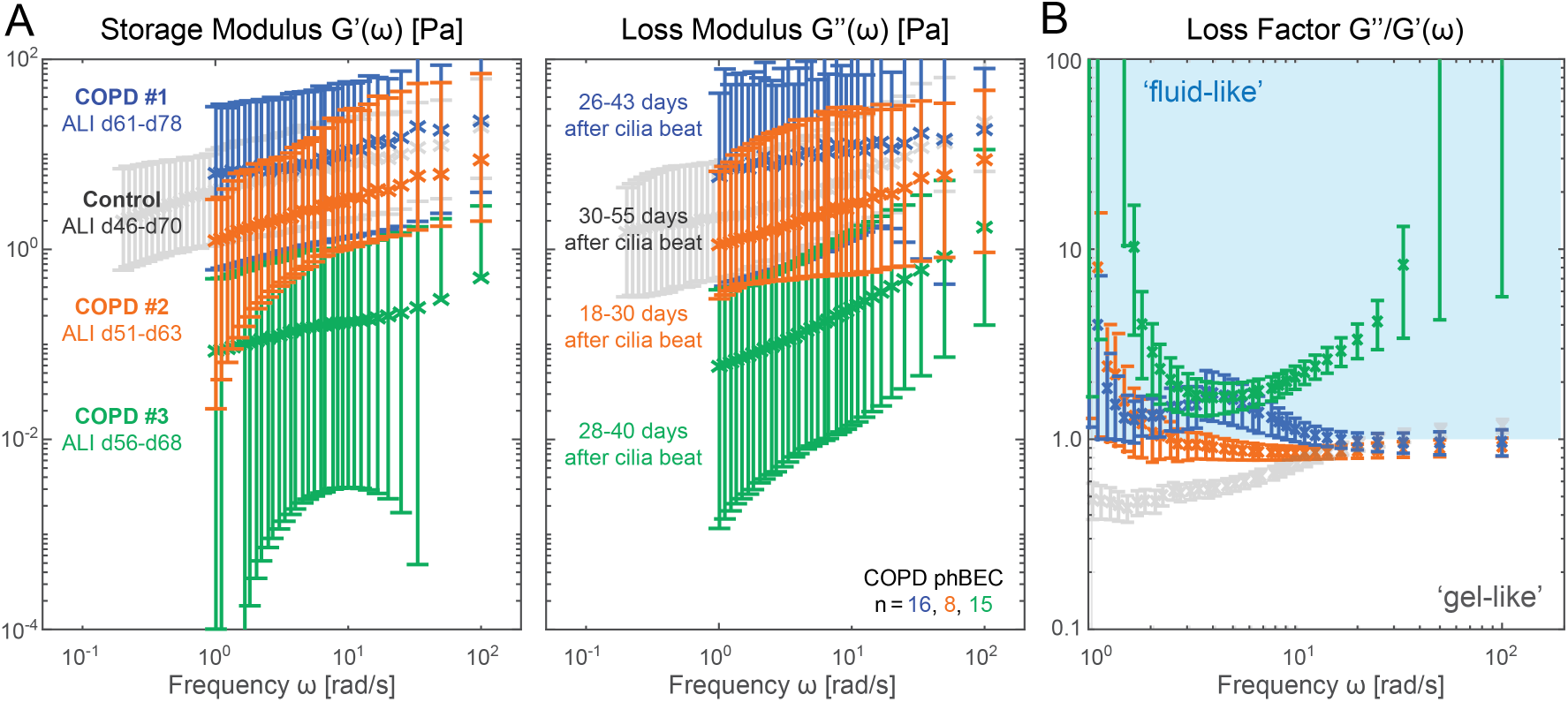
Donor-to-donor variability in mucus viscoelasticity from COPD ALI cultures. **A.** Storage and loss moduli of mucus extracted from three COPD phBEC donors differentiated in PC ALI medium (blue, orange, and green markers), compared with control PC cultures of similar ALI age (days 46 and 70 in Figure 3, light-gray markers). Markers show the median moduli. At each frequency, the lower and upper error-bar limits are the minimum 10th percentile and maximum 90th percentile, respectively, among the *q*-wise estimates from all recorded ROIs across the stated samples. **B.** Corresponding loss factors, *G^′′^/G^′^*. Each marker shows the mean across replicate measurements, and error bars indicate the standard error of the mean. Labels indicate ALI age and the number of days after visible ciliary beating was established across most of each insert. The analysis included *n* = 16, 8, and 15 samples from COPD donors #1, #2, and #3, respectively. Figure S5 (Supporting Information) shows donor-level heterogeneity at selected frequencies.

range, and donor #3 showed lower storage modulus and higher loss factors, consistent with a more fluid-like or diluted extraction (Figure 4A,B). Thus, despite similar clinical disease staging, COPD-derived cultures showed pronounced inter-donor variability, including partial overlap with healthy control cultures of comparable age. This pattern agrees with clinical sputum rheology studies reporting patient-specific variation and overlap between chronic airway disease and healthy-control ranges.^[33,96^ ^]^ Within this limited donor set, higher moduli were observed in cultures from donors with higher reported pack-years or more recent smoking cessation (Figure 4; **Table 1**), supporting the use of high-throughput mucus rheology as an additional quantitative metric for future COPD heterogeneity and disease-subtyping studies.^[52^ ^]^

**Table 1:** Donor information for primary airway epithelial cultures. Donor identifiers are reported as used throughout the manuscript and figures.

| **Commercial control donors** |  |  |  |  |  |  |
| --- | --- | --- | --- | --- | --- | --- |
| **Source** | **Cell type** | **Donor** | **Sex** | **Age** | **Ethnicity** | **Cause of death** |
| Cell Systems, FC-0016 | SAEC | 8938 | Female | 56 | Caucasian | Cerebrovascular accident / stroke |
| Cell Systems, FC-0035 | HBTEC | 7783 | Male | 12 | Caucasian | Self-inflicted gunshot wound |
| Cell Systems, FC-0035 | HBTEC | 9439 | Male | 48 | Asian | Cerebrovascular accident / stroke |
| **COPD donor-derived phBEC cultures** |  |  |  |  |  |  |
| **Source** | **Cell type** | **Donor** | **Sex** | **Age** | **Smoking history** | **COPD stage / clinical notes** |
| CPC-M bioArchive | phBEC | COPD #1 | Female | 64 | 80 pack-years; smoke-free for 4 years | COPD GOLD IV/E with emphysema and mild pre-capillary pulmonary hypertension |
| CPC-M bioArchive | phBEC | COPD #2 | Male | 63 | 30 pack-years; smoke-free for 8 years | COPD GOLD IV/D |
| CPC-M bioArchive | phBEC | COPD #3 | Female | 55 | 30 pack-years; smoke-free for 9 years | COPD GOLD IV/E with emphysema and pre-capillary pulmonary hypertension |

### 2.4 Rapid Phenotyping of In Vitro Disease Simulations

To show how our workflow can directly benefit translational research, we applied our measurement approach to three simulated disease conditions: (i) ALI culture exposed to cigarette smoke extracts, (ii) cystic fibrosis (CF) conditions rescued by drug treatment, and (iii) interleukin (IL)-13 induced asthma-like conditions.

In **Figure 5A-C**, we compared the rheology of diluted mucus collected from cigarette smoke extract exposed conditions (CS) with that from untreated cultures (UN). Both storage and loss moduli from the CS condition were slightly lower than those of the UN conditions, consistent with studies showing that human sputum in light smokers exhibits reduced viscosity.^[37,49^ ^]^ Importantly, we were able to calculate that normalized storage modulus decreased more than normalized loss modulus, suggesting that the CS exposed mucus was more fluid-like, possibly related to the onset of barrier function breakdown. Note that the loss factor of the untreated samples was below unity, unlike the previous PC day 70 control shown in Figure 3A, because mucus was allowed to accumulate on the apical surfaces as these cultures were never washed nor humidified until the day of extraction.

**Figure 5:**
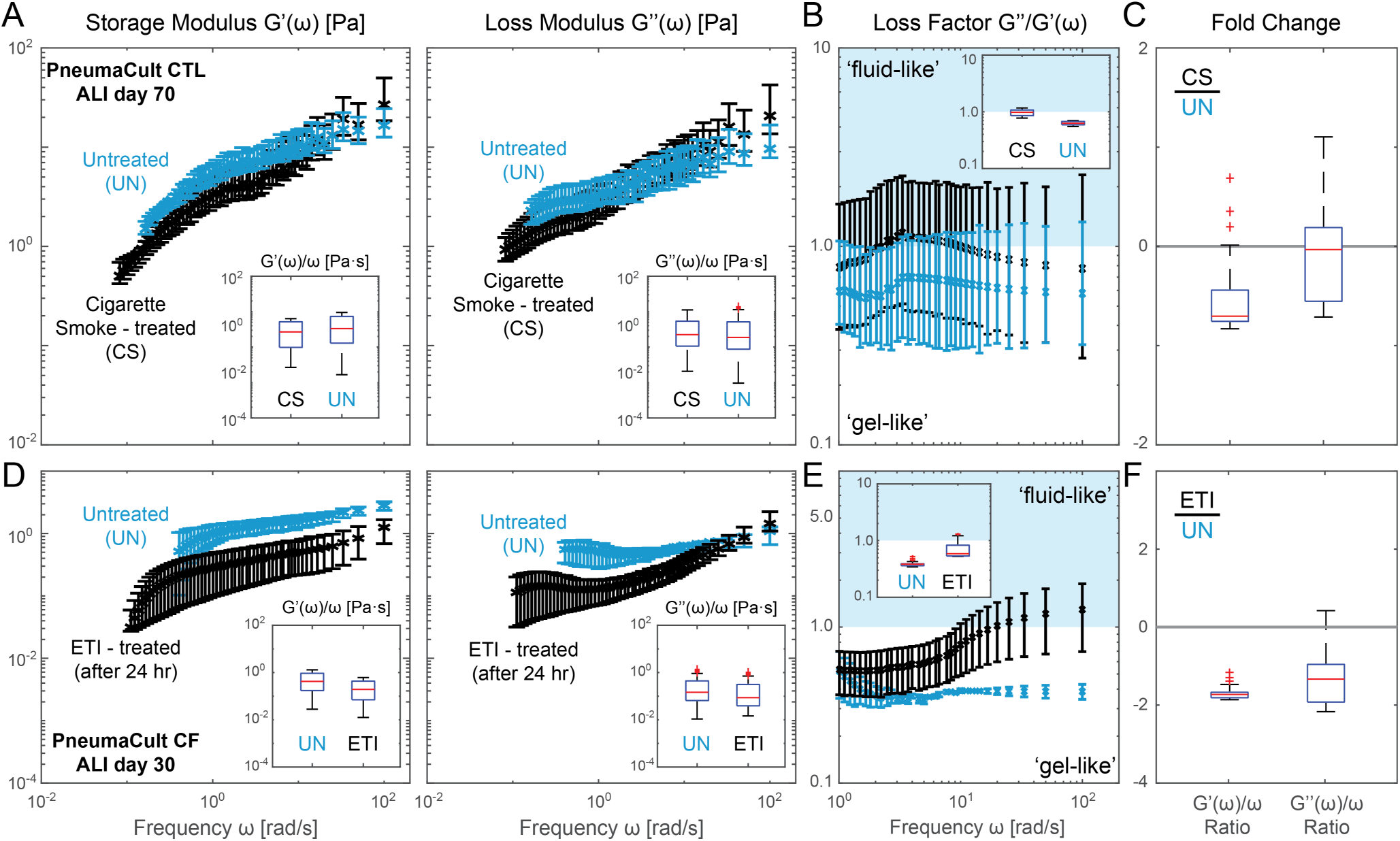
Rheological changes of *in vitro* airway mucus in response to apical treatments. A–B. Viscoelastic moduli and loss factors of mucus extracted from ALI cultures exposed to cigarette-smoke extract as a model of subchronic exposure (black, CS) or left untreated (blue, UN). A reduction in viscoelastic moduli and increase in loss factor is induced by cigarette smoke, hinting a potential compensation due to irritation. **C.** Fold change of the normalized storage (left) and loss (right) moduli between CS treated and untreated sample. **D–E.** Moduli and loss factors measured *in situ* in iPSC CF ALI cultures before (blue, UN) and after 24 h of ETI treatment (black, ETI). **F.** Fold change of the normalized storage (left) and loss (right) modulus from ETI treated inserts and untreated ones. In the modulus panels, markers show the median. At each frequency, the lower and upper error-bar limits are the minimum 10th percentile and maximum 90th percentile, respectively, among the *q*-wise estimates from all recorded ROIs across the stated technical replicates. Loss-factor markers show the mean across technical replicates, with error bars indicating the standard error of the mean. CS experiment was performed on mucus extracted 70 days after air lift from *n* = 2 technical replicates of a single donor at approximately 1:4 v/v dilution in PBS. CF experiment was performed *in situ* on *n* = 2 inserts from a single iPSC CF donor.

We further demonstrate the versatility of DDM rheology by performing in-situ measurement of iPSC-derived CF ALI cell cultures (iALI),^[97]^ where total amount of secreted mucus was too little to be extracted. Measurements were performed at day 30 after air lifting and minutes after ciliostasis induced by basal application of cinnamaldehyde;^[98,99^ ^]^ see Materials and Methods. Compared to UN, CF culture

exposed to 24 hours of Elexacaftor, Tezacaftor and Ivacaftor (ETI) treatment showed reduced moduli, increased loss factor, and near two-fold reduction in both normalized moduli; see Figure 5D-F. This finding is consistent with prior reports, hinting that ETI was able to partially correct mucus thickening and rescue the CF phenotype in the iALI cultures.^[20,34^ ^]^

Similar in-situ measurements were performed on pHAEC in IL-13 induced asthma-like inflammatory conditions.^[100–102^ ^]^ Previous studies suggested that these conditions promote a spatially heterogeneous mucus layer containing distinct domains dominated by different mucins.^[32^ ^]^ Hence, we were curious if IL-13 treatment would result in spatial heterogeneity of mucus rheology as well. Leveraging the positional information from in-situ techniques, we mapped viscoelastic moduli and loss factors as a function of distance from the insert center (encoded by marker darkness in **Figure 6**). In untreated controls, moduli from different regions clustered tightly, indicating uniform mechanical properties. By contrast, IL-13 treated mucus exhibited pronounced heterogeneity (compare Figure 6A and D), with localized stiffening near the insert center (darkest gray markers) but decreased viscosity in other regions. This pattern suggests that IL-13 drives spatially uneven mucus remodeling, likely reflecting a combination of local accumulation due to impaired mucociliary transport and mucus tethering, and barrier dysfunction with medium leakage under prolonged high-dose treatment schedule (100 ng·mL^−1^ for 32 days).^[32,101,103–105^ ^]^ Consistent with this interpretation, loss factors were overall elevated in IL-13 treated mucus (Figure 6B inset), but shifted toward more gel-like behavior in specific spatial regions and frequency ranges. Nor-

**Figure 6:**
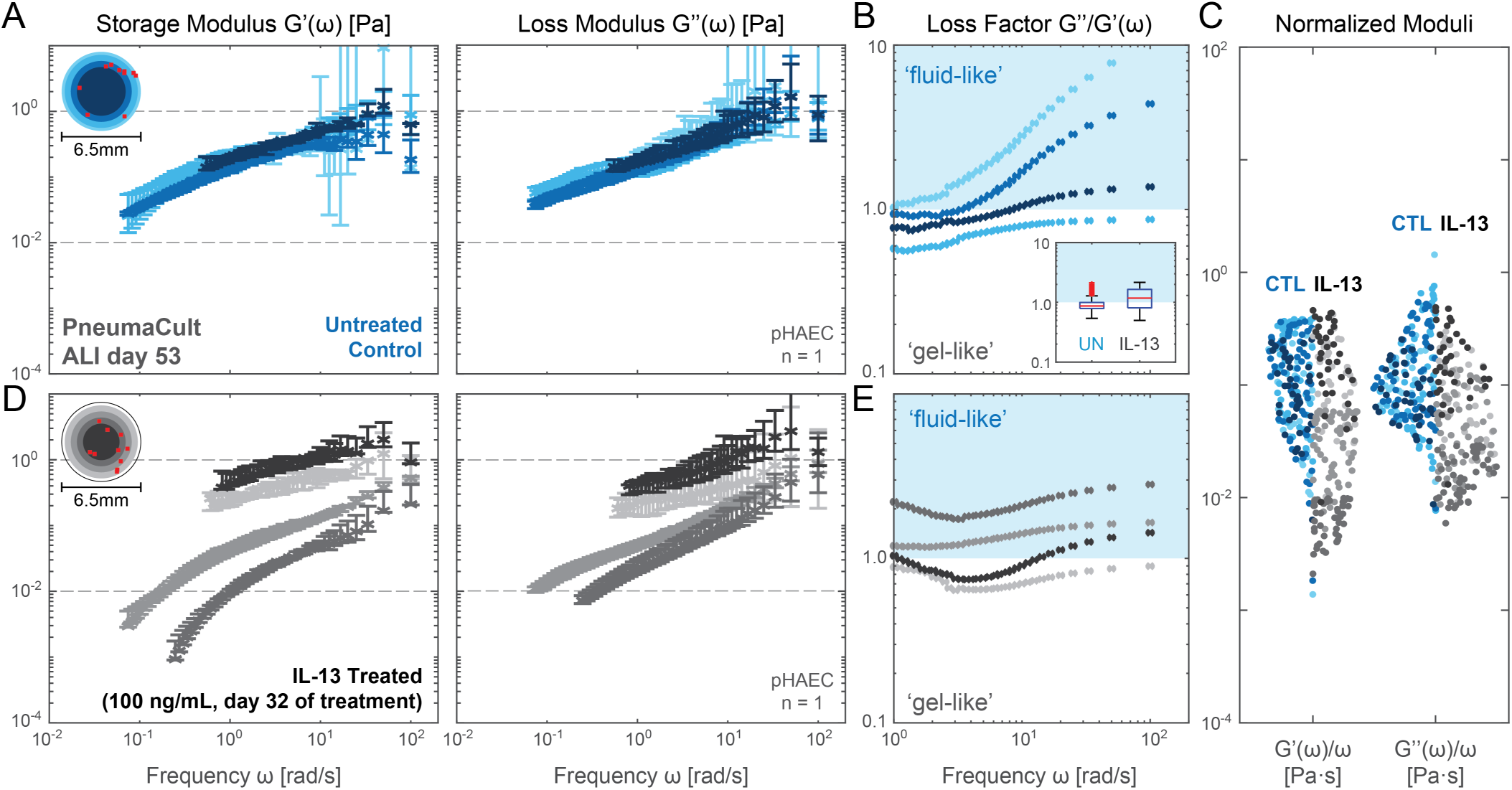
In-situ characterization of rheological heterogeneity from asthma-like in-vitro experiments. A and. **D.** Storage and loss moduli of mucus measured *in-situ* from untreated (blue tinted markers, CTL) and IL-13 enriched conditions (grayscale markers, IL-13). For each distance group and frequency, the modulus marker shows the median across valid ROI-level estimates. The lower and upper error-bar limits are the minimum *q*-wise 10th percentile and maximum *q*-wise 90th percentile, respectively, among the valid ROIs in that distance group. Marker shades are grouped based on the distance *r* from the center of ROI to the center of the insert in 0.5 mm intervals. From darkest to lightest, the control markers represent ROI taken at *r ≤* 2, 2 *< r ≤* 2.5, 2.5 *< r ≤* 3, or *r >* 3, respectively. Similarly, the IL-13 markers represent ROI taken at *r ≤* 1.5, 1.5 *< r ≤* 2, 2 *< r ≤* 2.5, or *r >* 2.5, respectively. For actual ROI locations, see red markers in top-left insets. The choice of distance groups was determined based on the actual location of the observed mucus droplets. In control condition, mucus migrated in bulk towards the edge of the insert as the result of racetrack-like global mucociliary clearance. In IL-13 condition, cells near the edge of insert experienced visible shrinkage and medium leakage. **B and E.** Loss factors computed from the median moduli for both conditions following the same marker definition. **C.** Scatter plot comparison of normalized storage and loss moduli between untreated and IL-13 conditions. Width of the scatter indicates kernel density estimation in log space. Measurements performed on *n* = 1 insert per condition using the same pHAEC donor.

malized moduli plots (Figure 6C) further showcase both the increase in heterogeneity and local stiffening in response to IL-13 treatment (note the upward shift from dark blue CTL markers to dark gray IL-13 markers).

In summary, we demonstrated that high-throughput DDM can detect subtle change of mucus state in response to different treatments and quantify mucus spatial heterogeneity in-situ in live ALI cultures. This rapid detection of scale-dependent changes in mucus properties is a crucial step toward disease-relevant modeling of mucus accumulation dynamics, mucus–cilia interactions, and functional alterations in mucociliary clearance.

### 2.5 Osmotic Swelling Assays from Rare Clinical Mucus Specimens

Not only can our approach process many microliter-sized samples for high-throughput studies, it also enables aliquot-based parallel measurements of scarce clinical mucus specimens. Since such specimens are often collected with buffers or processed in solutions, we aimed to determine how different submersion times alter the viscoelastic properties of the mucus samples in absence of aggressive homogenization or mixing. We used clinically-extracted cervical mucus at two time points of the menstrual cycle, when mucus is known to drastically differ in viscoelasticity,^[60,61^ ^]^ and tested how incubation in phosphate-buffered saline (PBS) buffer would distort these differences.

In **Figure 7A**, we verified that luteal phase cervical mucus (blue) indeed exhibited higher storage and loss moduli compared to the ovulatory phase mucus (red) from the same donor. After measuring the original fresh samples, technical replicates were created and submerged into PBS (pH 7, calcium-free) with standard pipettes at ∼ 1:100 v/v dilution ratio; see Materials and Methods. Care was taken to prevent droplet homogenization with PBS. Instead, mucus was left to swell gently via osmotic pressures.^[106–108^ ^]^ The PBS-exposed samples were then incubated in stationary condition at 4^◦^C for 5 minutes and up to one week before getting centrifuged at 400 RCF units for 10 minutes before extraction and measurement.

**Figure 7:**
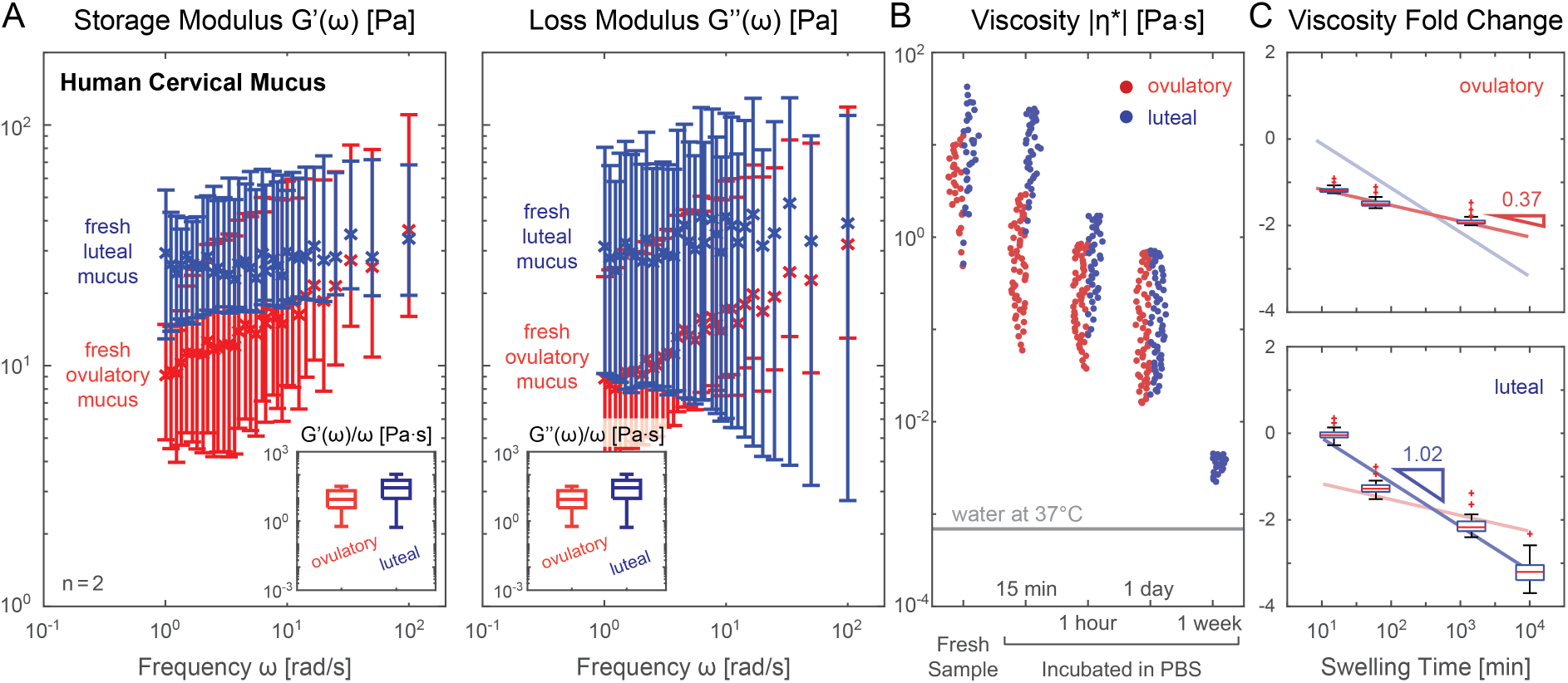
Characterization of viscoelastic change of clinical cervical mucus samples in response to osmotic swelling. **A.** Storage and loss moduli derived from clinically extracted cervical mucus at luteal (blue shaded markers) and ovulatory phase (red shaded markers), both from fresh collection (darker shade) and after exposure to PBS without homogenization (lighter shade). The luteal phase viscosity at higher frequencies are most likely outside the sensitivity limit of our methodology and thus discarded in later statistical analyses. Modulus markers show the median. At each frequency, the lower and upper error-bar limits are the minimum 10th percentile and maximum 90th percentile, respectively, among the *q*-wise estimates from all recorded ROIs across the stated technical replicates. **B and C.** Complex viscosity (B) and fold change of the luteal and ovulatory mucus before and after different PBS incubation (C). Power law fits based on the median of each fold change value and its power coefficient are also shown in (C). Measurements were performed on the same donor example with *n ≥* 1 technical replicates per swelling conditions.

Compared with pre-exposure samples, swelling dynamics as a function of incubation time were clearly reflected in the drop in absolute viscosity (Figure 7B-C). After 24 hours of incubation, the absolute viscosity of both mucus phases stabilized to similar distributions, and luteal-phase mucus appeared to homogenize naturally after 1 week of PBS exposure, as indicated by the collapse of its final viscosity distribution. The two distinct trajectories suggest that our ovulatory sample was more susceptible to osmotic swelling than the luteal-phase mucus, particularly under short-term exposures. This susceptibility contributed to the apparent steepening of the fold-change slope for luteal mucus swelling (Figure 7C) and could be a fluid-dynamic manifestation of the superior barrier function of luteal over ovulatory mucus. These assays also highlight how buffer exposure and incubation time can substantially alter mucus viscoelasticity, underscoring the need for controlled hydration, temperature, and storage conditions when processing mucus specimens. If applied at scale, our technology could be used to assess mucus rheology as part of standard routine across clinical specimens with diverse tissue origins.

## 3 Discussion

We present a novel general platform that enables high-throughput microrheology from microvolumes of mucus. Intriguing future applications in the clinic include the measurement of the currently disposed mucus fraction of nasal brushings and other cytology tests, which are gaining rapidly in popularity for precision medicine due their minimal invasiveness and amenability with multi-omics analysis.^[109–111^ ^]^ Assaying the mucus rheology in the same sample will add another personalized data point “for free.” Our platform also seamlessly integrates with state-of-the-art preclinical research by matching the throughput and mucus volumes of standard microtiter plates for cell culture while also enabling in situ measurements.

Importantly, this work shifts mucus rheology from a low-throughput, specialized measurement to a scalable and broadly accessible modality.

Our results establish that our optimized DDM microrheology workflow quantifies viscoelastic properties of microliter-scale mucus samples with minimal operator time. This efficiency enabled a single user to perform several dozen measurements per day, a throughput rarely attainable with tracking-based approaches. We also established standardized procedures for mucus extraction, handling, and storage across different culture sources and clinical samples. Clearly defining these often overlooked or poorly documented steps can reduce operator-to-operator variability and will be a key determinant of reproducibility, helping to minimize sources of variability that likely underlie inconsistencies between prior reports. By pairing a high-throughput assay with a rigorously standardized pipeline, our approach provides a practical route beyond sporadic measurements toward the large-scale datasets and longitudinal monitoring essential for understanding how mucus mechanics contribute to health and diseases of airway,^[112^ ^]^ reproductive tract,^[14^ ^]^ and other epithelial barriers. The COPD donor-derived airway cultures illustrate this need: despite similar clinical disease staging, they showed substantial variability in mucus rheology, consistent with the clinical and biological heterogeneity of COPD and with reported patient-specific differences in sputum viscoelasticity.^[33,96^ ^]^ Because in vitro culture can alter or amplify donor phenotypes, larger studies will be needed to determine how culture-derived mucus rheology relates to smoking history, clinical measures, subtype, or disease stage. Our platform provides the standardization and throughput required for such studies, while also supporting the maturation and integration of simulations of mucociliary clearance and airway surface liquid transport for disease modeling and drug delivery applications by relaxing the reliance on idealized constitutive laws and using experimentally grounded, condition-specific rheological inputs to enable physiologically relevant predictions for personalized medicine.^[113–1]^ Alternative technologies face significant throughput or sample volume trade-offs (**Figure 8**). Laser scattering methods such as diffusing wave or dynamic light scattering can provide extremely high-frequency bulk responses but require 12 to 150 µL samples and specialized devices.^[119–121^ ^]^ Optical tweezers and magnetic probe-based assays can access nonlinear behavior in-situ, but only at discrete points and come with substantial operational complexity.^[91,94,99,122–126^ ^]^ In contrast, our fluorescent-particles-based workflow is easier to handle and can capture wide fields of view that probe both in-sample and in-situ mucus heterogeneity. Commercial microfluidic viscometers such as Rheosense are limited to only measuring viscosity (i.e., not revealing elasticity) and require at least 19 µL per measurement to reach their maximum throughput of 96 assays in 24 hours.^[81^ ^]^ Finally, macro-rheometers optimized for mucus analysis such as Rheomuco require 20 µL of ALI mucus or 500 µL of sputum per test.^[33,127^ ^]^ Therefore, while these techniques remain indispensable for specialized queries, none are compatible with the routine, microliter-scale, high-throughput studies that our DDM-based approach makes feasible.

**Figure 8:**
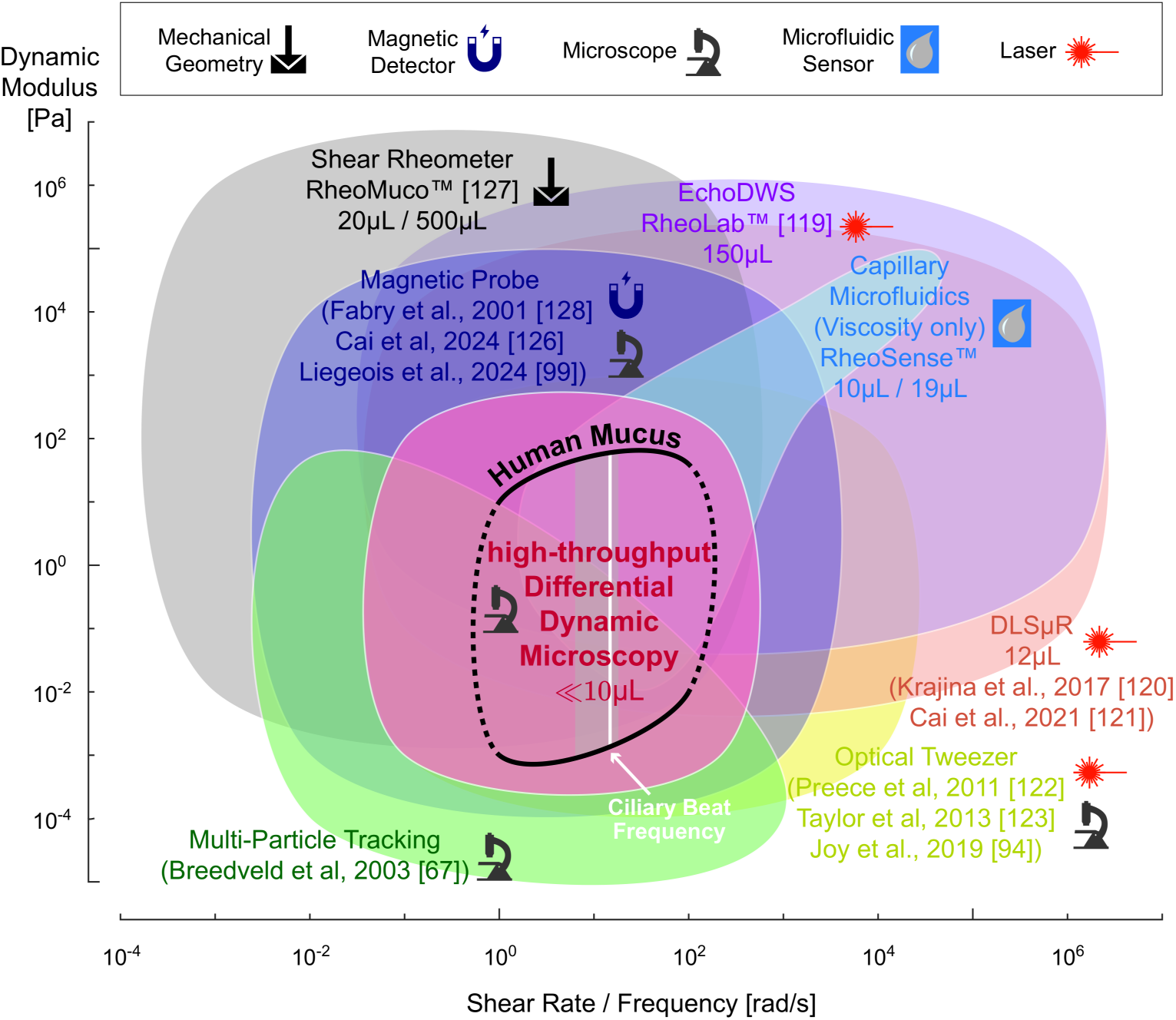
DDM is uniquely suitable for user-friendly, high-throughput assessment of low volume mucus samples. Accessible shear modulus and shear rate / frequency range of common rheology methods compared to human mucus modulus at frequencies relevant for biomedical research (black line).^[8,37]^ Ciliary beat frequency range based on our *in vitro* ALI culture measurements, with white line indicating median.^[30]^ Symbols indicate type of equipment required, and where applicable, we also indicate the required minimum sample volume for each method. Multi-Particle Tracking (light green) and Differential Dynamic Microscopy (pink) ranges are based on prior work and our reported setup.^[67]^ Dynamic Light Scattering (orange), Magnetic Probe (dark purple), and Optical Tweezer (yellow) ranges are based on published values.^[94,99,120–123,126,128]^ Mechanical shear rheometer (gray) values are based on our experience with Anton Paar devices. Sample volumes for RheoMuco device are the minimum ALI mucus volume reported in online FAQ and the recommended sputum volume from literature.^[127]^ Capillary microfluidic viscometer (light blue) and Diffusing Wave Spectroscopy (light purple) ranges are based on values reported by RheoSense m-VROC II and VROC Initium 1+ and LS Instruments RheoLab devices^[119]^, respectively.

Limitations of our workflow and refinements required for direct clinical use are as follows. While DDM is well suited for the quantification of linear viscoelasticity important for mucociliary clearance and particle permeability, it cannot directly capture strong nonlinear yield-stress responses that arise under cough-like shear. Clinical significance linking rheological readouts to clinical endpoints such as infection clearance, airflow obstruction, or drug response will also hinge on follow-up studies using larger donor cohorts to establish normative ranges beyond our proof-of-concept demonstrations. While our in-situ approach extends applicability to in vitro cultures that produce limited secretions, the transient inhibition of ciliary beating needed for these measurements can compromise long-term tissue integrity, especially in iPSC-derived cultures (see Supporting Information), limiting current reliable measurements to single time point per replicate. Finally, precautions ensuring stage mechanical stability, vibration isolation, and dust-free optics that are routine in research laboratories might require deliberate optimization in non-specialist point-of-care facilities to obtain reliable DDM readouts.^[80^ ^]^

Further improvements could extend the reach of this workflow without compromising its throughput. Open-source DDM implementations such as fastDDM can be readily integrated into our analysis pipeline to accelerate image-structure-function calculations, providing orders-of-magnitude reductions in processing time that might shrink *total* preparation and analysis time to within minutes per sample.^[80^ ^]^ Low-

cost imaging platforms, for example, webcam-based setup combined with deep-learning based frame interpolation and compensation for dropped frames^[129,130^ ^]^, could further improve accessibility in resource-imited laboratories and point-of-care settings. Because DDM preserves a direct correspondence between real-space information and reciprocal-space dynamics, in-sample spatial heterogeneity can be probed systematically by varying analysis window sizes, offering multi-scale insights particularly valuable for in-situ studies.^[131^ ^]^ Moreover, image sequences recorded for our one-point rheological analysis could also support two-point correlation analysis using optical-flow-based techniques^[132^ ^]^. This extension could yield robust measurements that better reflect bulk rheological response, provided that acquisition parameters such as tracer density, field-of-view size and recording duration are appropriately optimized. Taken together, continued advances in imaging hardware and computational methods should enable near-real-time DDM analysis to be integrated directly into microscopy platforms. Such an implementation would facilitate systematic testing of tracers with different sizes and surface chemistries across diverse mucus samples, thereby quantifying probe-dependent variations and establishing more robust criteria for tracer selection. In summary, by lowering the barrier to high-throughput mucus rheology, our workflow enables the correlation of biophysical mucus properties to clinical outcomes at an unprecedented scale. Knowing how rheological shifts link to environmental exposures or genetic predispositions could help identify new disease mechanisms, reveal biomarkers for muco-obstructive disease staging, and support disease subtyping, risk prediction, and the systematic evaluation of drug delivery and treatment strategies, ultimately advancing precision medicine approaches to mucosal barrier disorders.

## 4 Methods

### Study Design

This proof-of-concept study was designed to demonstrate the feasibility and throughput of our microrheology workflow across heterogeneous human mucus types, including airway mucus from donor-derived ALI cultures and cervical mucus from clinical samples. Sample size was determined by donor availability and culture success rates, with the intent of maximizing the number of technical replicates per condition. This design prioritized breadth and methodological validation by choosing experimental conditions with known changes to mucus rheology, rather than generating sufficient statistical power to probe unknown conditions, consistent with our objective of establishing general applicability. During experiments, samples were excluded when bubbles or leakage prevented localization of mucus in microscopy, or when unintended inclusion of beating cilia occurred. Cell-culture supernatants were also regularly screened for microbial contamination as well as mycoplasma using the MycoAlert Mycoplasma Detection Kit (Lonza, LT07-318) according to the manufacturer’s instructions. Other excluded datasets include reconstituted porcine MUC5AC solutions in neutral pH (used only for protocol development) and bulk mucus extraction from IL-13–treated and COPD donor ALI cultures in which barrier dysfunction caused significant medium leakage into the collected samples. No numerical outliers were removed during analysis. Technical replicates (small n) consisted of repeated measurements using mucus from the same donor and experimental condition, whereas biological replicates (big N) consisted of independent donor numbers or clinical samples. Power of the analysis was also enhanced via averaging over distinct regions of interest (ROIs) taken at different positions within each sample. Exact replicate information and sample origins are provided in relevant sections below. No predefined rules were applied to limit mucus extraction; data collection concluded when sample material or culture viability was exhausted. Randomization and blinding were not implemented, as the study focused on methodological development and relied on single-donor or limited-cohort sources. All data collection and analysis were performed by the study investigators. While exploratory in scope, this design establishes feasibility across multiple human mucus types and lays the groundwork for future multi-donor, statistically powered investigations. Donor characteristics of the primary airway epithelial cultures are summarized in Table 1.

### Mucus sample holders

Polidimethylsiloxane (PDMS) spacers (18 x 18 mm) were cut from 0.25 mm foil (MVQ Silicones, super clear) with 2–6 mm circular apertures, mounted on glass slides, and pre-seeded by evaporation drying of 1:2000 (v/v) suspensions of 500 nm yellow-green carboxylated polystyrene beads (Thermo Fisher Fluorosphere F8813) in the dark. After loading mucus samples, the chambers were sealed with #1.5H glass coverslips. A parallel Milli-Q-water-filled chamber was used to assess the post-desiccation effective bead hydrodynamic radius via the Stokes–Einstein relation. We found that the effective radius inflates from the nominal value of 250 nm to 280–300 nm on average at 37 ^◦^C; see Supporting Information Section 3.4 for more detailed analysis. The nominally 500 nm diameter carboxylated beads were selected to ensure sufficient coupling to the mucin network, so that the measured rheological response reflected the mucin gel rather than the solvent.^[8,62,91^ ^]^

### Reconstituted MUC5AC gels

Mucin (MUC5AC) was purified from porcine stomachs as described previously.^[85^ ^]^ The purified mucin was dissolved in de-ionized water at 4.4% w/v and kept overnight in a cold room to facilitate proper mucin solubilization. One hour before the shear rheometer measurements, 100 mm acetate buffer (pH 4) was added to suitable solutions of this mucin stock to obtain ca. 300 µL of 1,2,3, and 4% w/v mucin gels. 150 µL samples of those gels were used for shear rheometry; the remaining samples were stored at 4^◦^C until sealed into GeneFrame 25 µL chambers for DDM measurement within 12 hours. Test performed on a single sample with 4 − 15 ROIs recorded per sample.

### Medium condition ALI cultures

Control human primary airway (bronchial/tracheal and small airway) epithelial cells (donors 7783 and 8938; Table 1) were obtained from Cell Systems. Cells were supplied with vendor quality-control documentation indicating negative sterility testing for mycoplasma, bacterial and fungal contamination, and negative PCR testing for HIV-1, HIV-2, HBV and HCV. The first passage cells were expanded in collagen I coated 75 cm^2^ tissue culture dish in bronchial epithelial cell medium (BEpiCM) (ScienCell (Sanbio), SCC3211-b, Cat. No. 3211) until ∼90% confluency. Expanded cells were seeded on collagen IV (300 µg·mL^−1^) coated 6-well 0.4 pore diameter PET Transwell membranes (Corning, 3450) at a density of 500K cells per insert (∼135K cells/cm^2^). The cells were cultured in BEpiCM containing 1 nm EC23 (Tocris) until fully confluent. Primary human bronchial epithelial cells (phBECs) from COPD patients were obtained from the CPC-M bioArchive at the Comprehensive Pneumology Center Munich (CPC-M; Table 1).^[95^ ^]^ The first passage cells were expanded until 90% confluent in a custom hydrogel coating (ECMmix) on 100 × 15 mm tissue culture treated dish (Santa Cruz, sc-200286). The custom coating solution composed of 10 mL PBS at 4^◦^C, 100 µL PureCol (3 mg·mL^−1^, Cellsystems, Cat. No. 5005-100ML, Lot 8201), 100 µL BSA (0.1%), and 50 µL human fibronectin (1 mg·mL^−1^, Promocell, Cat. No. C43060). The coating solution was vortexed and then incubated inside the dish in incubator at 37^◦^C for at least 2 hours. After incubation, the solution was aspirated and 10 mL of BEpiCM was added to equilibrate in the dish before cells were added. Expanded COPD cells were seeded on collagen IV (300 µg·mL^−1^) coated 12-well 0.4 pore diameter PET Transwell membranes (Corning, 3401) at a density of 150K cells per insert (∼135K cells/cm^2^). The cells were cultured in BEpiCM containing 1 nm EC23 (Tocris) until fully confluent. Once all tissues were confluent, differentiation was induced by introducing air liquid interface (ALI) via removal of the apical medium (day 0 of ALI culture) and the use of PneumaCult ALI (STEMCELL Technologies, PC) or BEpiCM:DMEM 50:50 + 1 nm Ec23 medium (BEGM-based) supplied through basal chamber. During routine maintenance (every two days), cultures were inspected using a ZEISS Primovert inverted cell culture microscope under a 10×objective to assess the existence of global ciliary beating motion. The first observed date was recorded as the point of ciliogenesis. This criterion corresponds to the stage when ciliary patches are extensive enough and the ciliary motion is sufficiently mature.

### Cigarette smoke ALI culture and treatment

Human primary airway (bronchial) epithelial cells (pHAECs) were acquired from cancer-free smokers at the University of New Mexico through diagnostic bronchoscopy and were stored in a de-identified manner. Donor culture plates were coated with lab-made 804G medium using RPMI (Gibco, 11875119) + 10% FBS (Gibco, A5256801), and SABM with SAGM bullet kits (Lonza, CC-3118). At 80% confluency, cells were seeded onto 804G media-coated 12-well Transwell inserts and cultured in PneumaCult ALI (STEMCELL Technologies) until they were confluent. Once the tissues reached 100% confluency, differentiation was induced by removing the apical medium to establish an air liquid interface (ALI) setting (day 0 of ALI culture) and the use of PneumaCult medium supplied through basal chamber. Cigarette Smoke Extract (CSE) solutions were prepared from research cigarettes (3R4F, Center for Tobacco Reference Products, Kentucky Tobacco Research & Development Center, Lexington, KY) as previously reported, with additional methodological detail provided elsewhere.^[133,134^ ^]^ CSE treatment was performed by exposing cells for 1 h twice per week for 8 weeks either with vehicle (culture medium; UN condition) or medium containing 40 µg·mL^−1^ CSE (CS condition) from the basal side.

### IL-13 ALI culture and treatment

Human primary airway (bronchial/tracheal) epithelial cells (donor 9439; Table 1) were obtained from Cell Systems. Cells were supplied with vendor quality-control documentation indicating negative sterility testing for mycoplasma, bacterial and fungal contamination, and negative PCR testing for HIV-1, HIV-2, HBV and HCV. The first passage cells were expanded in collagen I coated 75 cm^2^ tissue culture flasks in bronchial epithelial cell medium (BEpiCM) (ScienCell (Sanbio), SCC3211-b) until ∼90% confluency. Expanded cells were seeded on ECMmix coated 24-well 0.4 pore diameter PET Transwell membranes (cellQART) at a density of 75K cells per insert (∼135K cells/cm^2^). The cells were cultured in BEpiCM until fully confluent. Once the tissues were confluent, differentiation was induced by introducing air liquid interface (ALI) via removal of the apical medium (day 0 of ALI culture) and the use of PneumaCult ALI (STEMCELL Technologies) supplied through basal chamber. Starting on day 21 of ALI culture, 100 ng·mL^−1^ of IL-13 treatment is added to the basal medium until measurement day.

### iALI CF culture and ETI treatment

A hiPSC line (MHHi002-A) carrying a homozygous CFTR Phe508del mutation as a diseased reference was used for measurements. As detailed previously, induced pluripotent stem cells (iPSCs) were cultivated on Geltrex in E8 media before differentiation, and then first differentiated towards definitive endoderm (DE) through STEMdiff Definitive Endoderm Kit (Stem Cell Technologies, TeSR-E8 Optimized).^[97^ ^]^ DE cells were further differentiated towards anterior foregut endoderm (AFE) by (i) 1-day treatment with 3 µm dorsomorphin (Merck), 10 µm SB431542 (provided by A. Kirschning, Leibniz University Hannover) and ROCK Inhibitor Y27632 (Tocris), and (ii) 1-day treatment with 2 µm IWP2 (Tocris) and 10 µm SB431542. AFE cells were differentiated towards NKX2.1 positive early lung progenitors via 8-day treatment of 10 µm BMP4 (R&D), 3 µm CHIR99021 (provided by A. Kirschning, Leibniz University Hannover) and 10 nm FGF10 (R&D). On day 13 of differentiation, NKX2.1 positive cells were enriched by applying Magnetic Activated Cell Sorting for the surface marker Carboxypeptidase M (CPM) using an anti-CPM antibody (FUJIFILM Wako). Enriched progenitor cells were seeded to 12-mm transwell (Corning) and cultured in small airway epithelial growth medium (Promo Cell) supplemented with 0.5 µm A–83–01 (Tocris), 0.5 µm DMH–1 (Tocris) and 5 µm ROCK Inhibitor Y27632 for four days. Afterwards, cells were treated with PneumaCult-ALI medium (Stem Cell Technologies) and airlifted after eleven days. All iALI cultures were matured for a total of 6 weeks and treated with 5 ng·mL^−1^ IL-4 (Peprotech) supplement until measurement. 14 days before measurement, cell cultures were washed apically with PBS and ciliary beat frequency were measured to ensure proper health and maturation. One day (> 24 hour) before measurements, ETI treatment composed of 3 µm Elexacaftor (Selleckchem), 18 µm Tezacaftor (Selleckchem), and 1 µm Ivacaftor (Selleckchem) was applied to half of the matured inserts for comparison.

### Medium comparison and COPD mucus extractions

Forty-eight hours before collection, the apical side of Transwells was incubated with 500 µL of Ca^2+^ infused PBS for 10 minutes (inside 37^◦^C incubator). Afterwards, the PBS was removed and 50 µL/cm^2^ of medium was added (*e.g.*, 233.5 µL per insert for 6-well plates, Corning 3450) to humidify the culture.^[135^ ^]^ Mucus was collected and stored at +4^◦^C in Eppendorf tubes sealed with Parafilm on days 20, 25, 27 (week 4), 46 (week 6), and 70 (week 10) of ALI culture until measurement day (as soon as possible and less than one work week). COPD cultures were measured from collections 51 and 63 days after airlifting, also with a humidifying medium added two days prior to measurement similar to the controls. The time from the day of ciliogenesis is also recorded on measurement day. In order to verify mucus storage could not significantly alter our measured outputs, a week-long storage test on two donors was performed; see Figure S2 (Supporting Information). Test performed on N=2 donors for control (7783 and 8938) and N=3 donors for COPD (COPD #1-#3) from n=2 or 3 inserts with 5-10 ROIs.

### Accumulated mucus extracted with paper device

To efficiently extract high-viscosity mucus from 6.5 mm transwell inserts, weighing paper (Whatman weighing boats and paper Z134112 WHA10347671) was cut into ∼ 3 mm wide strips, each folded in a zig-zag fashion with ∼ 5 mm spacings. Prepared paper strips were then put inside UV curing chambers 30 minutes per side for sterilization. To extract mucus droplets, the end of the paper strip was pushed into the transwell inserts with care inside the biosafety cabinet; see Video S1 (Supporting Information). Based on our testing, the folding geometry of the paper could absorb enough of the vertical applied pressure and kept cell layers intact as long as the side edges were kept away from the cell surface. Within a few seconds of contact, high viscosity mucus droplets were absorbed onto the paper strip. The paper was then taken outside of the biosafety cabinet and the attached mucus droplet was transferred into the prepared capillary chamber with a metal laboratory spatula as soon as possible to minimize dehydration. Test performed on N=1 donor (8938) from n=2 technical replicates with 8 ROIs per sample.

### Cigarette smoke treated mucus extractions

Cultures were not humidified nor washed on the apical side for the entire ALI culture duration (70 days, both CS and UN). On collection day, 30 µL of PBS (Calcium-free) was added to the apical side of the transwell (≈ 27 µL/cm^2^) for 3 minutes, and mucus was harvested using a positive displacement pipette (Gilson Microman E). Mucus samples were sent by overnight mail and kept at 4^◦^C until measurement day in the same week. A sample volume of 8 µL was first loaded in a 6 mm capillary chamber for imaging (results are outside the sensitivity capability of DDM), then the same loaded sample was diluted (mixed until homogenization) with 10 µL of Milli-Q water twice to enhance the measurable difference between CS and UN samples. This gives two final technical replicates with equivalent dilution ratio of 16/81≈1:4 v/v (results of Figure 5A-C). Dilution was conducted to enhance the difference between measured viscoelastic moduli between CS and UN samples; prior to dilution the results were outside the sensitivity zone of our method and yielded virtually indistinguishable results. Test performed on N=1 donor from n=2 technical replicates with 5 ROIs per condition.

### In-situ imaging

Carboxylated fluorescent beads (Thermofisher, FluoSpheres F8821, 1 µm diameter) were used for improved real-space visibility during in-situ imaging. They were first mixed at 1:5000 v/v dilution into culture medium and applied on the apical side of *target and control* inserts at 50 µL/cm^2^ volume *two days* before measurement to minimize undesired hydration effects. During measurements, PneumaCult culture medium enriched with 10–30 mm of cinnamaldehyde (CA) was put to the basal chamber of the transwell plate to inhibit ciliary motion. Note that since undissolved cinnamaldehyde reacts with polystyrene, care was taken to avoid polystyrene containers during preparation and optical artifacts during imaging. Once CA is applied, long distance objectives (WD 7.9mm at NA 0.4 instead of WD 0.55mm at NA 0.8) at the same 20x magnification on the same Axio Observer set-up were used to monitor ciliary beat frequency in-situ until no ciliary motion can be observed, whence DDM recordings were started. For induced pluripotent stem cell (iPSC) cultures (Figure 5D-F), 15 to 20 minutes of 10 mm CA treatment was sufficient to stop ciliary motion. For primary human airway epithelial cells (pHAEC) cultures (Figure 6), the time to full ciliostasis varied between donors and culture conditions and took anywhere from 15 to 60 minutes. During DDM recordings, care was taken to make sure the focal plane is around one cilia length above the cell surface to avoid imaging tracers not trapped inside the mucus layer. After imaging, basal chambers were washed with PBS and replenished with cinnamaldehyde-free medium immediately to minimize long-term toxicity effects. CBF were continuously monitored at regular intervals after imaging. For the CF in-situ test, n=2 inserts from N=1 donor was used with 6 ROIs per condition. For the IL-13 in-situ test, n=1 insert from N=1 donor (9439) was used per condition. A total of 10 ROIs were taken and they were each analyzed as four groups based on their distance away from the center of the insert as marked in Figure 6A and D.

### Cervical mucus collection and swelling

Cervical mucus samples were collected at the Department of Gynecology from Klinikum Rechts der Isar (MRI) via endocervical sampler cytobrush biopsy (Cooper-Surgical Inc.). All participants or their legal representatives provided informed consent. Samples were kept on the brush and stored in Falcon tubes at 4^◦^C (on ice during transport) and 10-11 µL-sized droplets were transferred with positive displacement pipette to be measured on the same day. For mucus swelling test, another ∼ 11 µL mucus sample was taken from nearby positions on the brush and lightly mixed with 830 µL of PBS (aliquot refrigerated with collected mucus) solution by pipette (1 mL) inside an Eppendorf tube. Care was taken such that the mucus droplets remained integral (one single visible blob) during the mixing process. Afterwards, 276 µL of PBS was added again and the Eppendorf tube was incubated for 5 minutes at 4^◦^C. Next, the Eppendorf tube was centrifuged at 400 RCF for 10 minutes at 4^◦^C. The mucus droplet was extracted from the supernatant fluid and placed into the capillary chamber for viscoelasticity measurements. We did not use the same sample for different swelling tests to avoid transfer losses. Test performed on N=1 donor with n=3 technical replicates for fresh ovulatory mucus, n=2 replicates for fresh luteal mucus, and single n=1 sample at different incubation times. 5 ROIs were taken for each sample.

### Shear rheometry

Small amplitude oscillatory shear measurements were performed using a commercial shear rheometer (MCR 302, Anton Paar, Graz, Austria) with a plate/plate geometry (bottom plate: P-PTD 200/AIR, Anton Paar; 25 mm steel measuring head: PP25, 79044, Anton Paar) and a plate separation of 300 µm.^[84,136^ ^]^ Pre-measurements were conducted in a stress-controlled manner at a torque of 0.5 µNm to ensure the characterization of linear viscoelastic responses. Then, frequency-dependent measurements were conducted in strain-controlled mode using the 1.5-fold value of the strain determined in the pre-measurements. Each 150 µL sample was measured 3 times over a frequency range of 0.01 to 10 Hz, with the frequency sweep going from maximum to minimum, vice versa and back again. Special care was taken so that the sample completely filled the gap between the two opposing plates. The instrument was outfitted with a humidity trap to minimize evaporation during testing. All measurements performed at 37^◦^C.

### Multi-particle tracking analysis

MPT is performed on the DDM videos using FIJI Trackmate with Differences of Gaussian (DoG) algorithm and simple Linear Assignment Problem (LAP) tracker.^[137^ ^]^ Manual thresholds were used to filter out tracks with low tracking quality, abnormally high mean intensity, short track lengths, and out of distribution mean track speed manually based on visual inspection. Before initiating Trackmate, video contrast is adjusted to contain 0.01% saturated pixels based on full stack histogram. To validate our high-throughput pipeline is performing on par with MPT and brightfield DDM, we performed measurements on homogeneous polymer solutions of 0 to 4% (w/w) Poly(ethylene oxide) solution in Milli-Q water; see Figure S4 (Supporting Information).

### Statistical Methods

All box plots visualize the median and 25th to 75th percentile as the box, with outliers (red crosses) outside of the whiskers defined to be more than 1.5 times the 25% to 75% range away from the box in linear units. Scatter violin plots show original data point spread based on kernel density estimates reconstructed with built-in MATLAB function KSDENSITY. In storage- and loss-modulus plots, markers show the median. At each frequency, the lower and upper error-bar limits are the minimum 10th percentile and maximum 90th percentile, respectively, among the *q*-wise estimates from all recorded ROIs across the stated technical replicates. These ranges are descriptive rather than fully propagated confidence intervals; sensitivity of *B*(*q*) estimation and tracer calibration are detailed in Supporting Information Sections 2.2 and 3.4, and the uncertainty hierarchy is summarized in Section 4. Normalized moduli and loss factors were calculated from the median of viscoelastic moduli at each measured frequency. In loss-factor plots, markers show the mean across technical replicates, and error bars indicate the standard error of the mean. In cases where only a single technical replicate was tested, error bars were omitted.

## Supporting Information

Supporting Information is available from the Wiley Online Library or from the author.

## Acknowledgements

We sincerely thank Prof. Eva Kanso for initial discussions on utilizing DDM for mucus rheology measurements. We are indebted to Dr. Wilhelm Bertrams and Prof. Mareike Lehmann for their valuable advice and numerous amounts of mucus extractions that helped us verify our throughput capacities and finalize the mucus collection protocols. We gratefully acknowledge the provision of human biomaterial (phBECs) and clinical data from the CPC-M bioArchive and its partners at the Asklepios Biobank Gauting, the LMU Hospital and the Ludwig-Maximilians-Universität München. We thank the patients and their families for their support. We are also thankful for Laurien Czichon and Nils Natrup for their invaluable help during the production of the iALI cultures. Lastly, we are grateful to Dr. Vandana Kaushal, Prof. Sylvester Holt and Prof. Thomas Hummel for coordinating a preliminary pilot study using nasal mucus samples that further strengthened our confidence in the workflow.

The authors acknowledge the use of OpenAI’s GPT5 models to support formatting and editing. All AI-generated suggestions were reviewed, revised, and approved by the authors, who take full responsibility for the accuracy and integrity of the work.

## Author Contributions

F.L. and J.C.N. designed the experimental pipeline, performed data analysis, and interpreted the results. A.T.S., D.R., F.L., A.P., N.T., R.A., and B.Z. optimized *in vitro* pHAEC culture and mucus collection protocols. B.M.N. and O.L. prepared reconstituted mucus samples and executed shear rheometry experiments. S.A. provided guidance for DDM methodologies. A.S.V. and Y.T. established cigarette smoke-exposed HBEC models and collected mucus samples. M.C.K. and R.O. established iPSC cystic fibrosis cultures and provided treatment protocols. E.E. and M.K. procured human cervical mucus samples and ensured compliance with ethical guidelines. M.G.S. and P.M. sourced COPD patient donor cell materials. All authors contributed to manuscript writing and approved the final version.

## Data Availability Statement

MATLAB code for the analysis pipeline and processed rheology measurement results supporting this study are available in the Zenodo repository at https://zenodo.org/records/17651732. Raw microscopy videos and measurement data that support the findings of this study are available from the corresponding author upon reasonable request, subject to privacy and ethical restrictions.

## Ethics Approval Statement

COPD patient-derived material collection was approved by the local ethics committee of Ludwig-Maximilians University of Munich, Germany (ethics vote No. 19-630). Cervical mucus material collection was performed in line with the principles of the Declaration of Helsinki, with approval granted by the Ethics Committee of the Technical University of Munich (Date 25.04.2022 / No. 2022-156-S-KH). Written informed consent was obtained for all study participants.

