## Supporting Information for "A general high-throughput mucus microrheology platform for quantitative and scalable mucus phenotyping"

### 1 Preparation and imaging protocols

#### 1.1 Chamber preparation

To prevent sample dehydration and facilitate its storage and reuse, we created custom capillary chambers from commercially available sheets of polydimethylsiloxane (PDMS) foil with 0.25 mm nominal thickness (Super clear, MVQ Silicones GmbH). This thickness was chosen to be compatible with the maximum working distance of our microscope objective. Using scissors and biopsy punches or a desktop vinyl cutter (Roland DG), 18×18 mm squares were cut from the silicone foil, with a 3 or 6 mm diameter hole centered in the middle in batch. These geometries yield a final measured holding volume of ca. 3 or 8 $\mu L$, respectively. For samples available in larger volume ($\geq 30$ $\mu$L), commercially available GeneFrame 25 $\mu$L (Thermo Fisher Scientific, AB0576) were used to mount the sample. Holder with smaller 1 or 2 mm diameter hole were also prepared in case the amount of collected mucus were insufficient to fill the 3 mm diameter chambers.

#### 1.2 Tracer preparation and calibration

After mounting the spacer to a standard microscope slide (VWR, 631-1553), we filled the chamber with a dilute solution of fluorescent tracer beads and allowed them to desiccate at room temperature in a light-tight box for 30–45 minutes. We used 500 nm diameter yellow-green carboxylated polystyrene beads (Thermo Fisher Scientific, Fluorosphere F8813) at 1:2000 v/v dilution in Milli-Q water (membraPure water purifier). Polystyrene particles with carboxyl surface chemistry and radii above 250 nm are chosen based on the expectation that they would be sterically hindered in mucin networks,[1–3]

Once all chambers had dried, one chamber was immediately refilled with Milli-Q water and sealed with a #1.5H glass coverslip for hydrodynamic-radius measurement. Specifically, the known viscosity of water $\eta_o$ was used to infer the hydrodynamic radius $r$ of the desiccated beads from the Stokes–Einstein relation $r = k_B T/(6\pi\eta_o D)$. Here, $k_B$ is the Boltzmann's constant, $T$ the measurement temperature, and $D$ is the diffusion coefficient that is equal to the slope of mean-squared displacement divided by 4; also see (4) and (5). Three accepted high-frame-rate water-control trials each comprised three raw FOV movies; see Figure S3. All three trials reached approximately 498 frames $\mathrm{s}^{-1}$ and contributed 11 frequency points to the primary 30–100 [rad $\mathrm{s}^{-1}$] window. In Trial #1, Trial #2, and Trial #3 order, their median loss viscosities before calibration were 0.6808, 0.6235, and 0.7013 mPa s, compared with the 0.6913 mPa s reference value for water at 37 °C. Inverting these values gave median effective radii of 289.5, 297.6, and 298.2 nm and a three-trial mean of 295.1 nm. The corresponding median $q$-wise radius bands were 282.7–293.0, 288.7–327.9, and 287.5–305.2 nm. The frequency-wise coefficients of variation within the selected window were 0.49%, 1.39%, and 0.37%, respectively. Finally, each trial's saved $G''/\omega$ values were multiplied by a single factor without frequency-specific correction $r_{\mathrm{input}}/r_{\mathrm{eff,trial}}$.

A companion water-filled chamber can also be prepared and imaged alongside each experimental batch to minimize batch-wise uncertainties due to minute changes in desiccation time and environment.

#### 1.3 Sample preparation

The mucus sample was introduced into the chamber lined with dried tracers using a positive displacement pipette (Gilson Microman E). To reduce bubble formation, the mucus was dispensed slowly and the pipette was set to hold 2 $\mu$L more than sample volume when filling the chamber, avoiding the introduction of air at the end of loading. If large bubbles did form, it was often possible to pop or remove the bubble by dragging it onto the PDMS spacer or poking it with the corner of a clean glass cover slip.

In order to ensure that a sufficient amount of dried tracer was present in the bulk of the sample, a sanitized small metal laboratory spatula was used to gently agitate the sample inside the chamber. In most cases, moving the spatula in both clockwise and counterclockwise directions for 10 to 30 seconds was sufficient to achieve adequate mixing.

Finally, a 1.5H glass cover slip was gently pushed onto the chamber starting from one edge to avoid bubbles and cracking. Once sealed, the prepared slides were stored at 4°C for later experiments, or placed for imaging into the temperature-controlled incubation chamber of the microscope preheated at 37°C. We ensured that the temperature control on the microscope reached equilibrium before imaging, since temperature fluctuations result in changes to Brownian motion and thus erroneous viscosity readout. Therefore, in all experiments, we waited at least 30 minutes after the incubator temperature sensor reported the desired temperature.

#### 1.4 Fluorescence imaging

We recorded all videos using an inverted epifluorescence microscope (ZEISS Axio Observer) equipped with a high-speed camera (Hamamatsu Photonics Orca Flash 4.0) and a 20x objective (0.8 NA, 0.33 µm·px$^{-1}$). Since DDM allows working with tracers smaller than the real-space optical resolution of the microscope, a 10x objective (0.3 NA, 0.65 µm·px$^{-1}$) was also tested, achieving good results. However, in order to better compare tracking-based analysis with DDM, we opted for the 20x objective with higher resolution, allowing individual tracer to be clearly resolved in space.

We first checked the sample in the bright-field channel to verify that no beating ciliated cells were present, as these can be accidentally removed from the tissue culture together with mucus. We noted down the location of microscopic bubbles and large aggregate/debris to be avoided. We found that for high viscosity mucus samples, minuscule amounts of drift ($< 0.1$ µm·s$^{-1}$) were sometimes present, potentially due to slow relaxation of mucus after slide movement, unevenly placed cover slip, and/or streaming around a distant bubble. While small amounts of unidirectional drift can be corrected during analysis, we re-sealed problematic samples to reduce drift, whenever possible. Finally, to minimize boundary effects, we measured Brownian motion in the central z-plane of the sample. This focal plane was computed by averaging the two most extreme z-values each representing beads located at the floor and ceiling of each chamber, using the maximum field of view available.

For video acquisition, we used the FITC/GFP channel. Most mucus samples were recorded for 20 s at 10 ms per frame (100 fps) and 30% LED power. For rapidly decorrelating, low-viscosity samples, we used minimum lags of 1–2 ms (500–1000 fps, depending on camera mode) and 100% LED power. The accepted water-control trials reported in Section 3.4 were recorded at approximately 498 fps. The 20 second video option was used whenever tracers were observed to vibrate back and forth around an equilibrium position in real time. The exposure values were chosen to maximize signal-to-noise ratio and minimize the global decay of fluorescent signals over time. Streaming option was always selected during time series recording, so the final video frame rate was determined directly by the exposure time setting. The nominal minimum lag was therefore approximately 10 ms for most mucus experiments and 1–2 ms for rapidly decorrelating controls. These acquisition values were not treated as universal validity cutoffs. Accepted lags were restricted during analysis by the physical condition $0 < x < 1$, the selected $q$ range, the availability or stability of the estimates of $B(q)$ and $D(q, \infty)$, and the removal of high- or low-lag artifacts. Depending on the heterogeneity of the sample, we recorded 3 to 10 ROIs, each at 256×256 pixels without binning at 0.33 µm·px$^{-1}$, while staying in the same z-plane and avoiding clusters of aggregated tracers. In our experience, a few aggregated beads did not cause significant problems, assuming most of the motion comes from isolated tracers. The nature of DDM analysis also means that insufficient contrast due to static background is not a concern, unlike in MPT.

After imaging, the entire slide was sealed with Parafilm for long-term storage. Our *in vitro* samples did not require additional antibiotics to prevent contamination, and viscosity values remained stable for one week after storage at 4°C; see Figure S2.

### 2 Differential Dynamic Microscopy

Viscoelastic moduli can be calculated automatically without user intervention from the recorded videos. Our analysis was performed in MATLAB on an Intel i9 computer with 128 GB of random access memory. The measured analysis time reported in the main text refers to this implementation. Below we outline the mathematical foundation of this calculation and highlight our custom adaptations from common practice that are necessary for high-throughput applications. Detailed derivations of equations (1) to (5) are available in prior references.[4–8]

#### 2.1 Image structure function.

The azimuthally averaged image structure function $D(q, \Delta t)$ is obtained as a function of the azimuthal wave number $q$ and time lag $\Delta t$ from the video frame sequences via

$$D(q, \Delta t) = \left\langle |\mathrm{FFT}[I(x, y, t + \Delta t) - I(x, y, t)]|^2 \right\rangle_{t,q}, \tag{1}$$

where $I(x, y, t)$ is the image intensity function over spatial coordinates $x, y$ and time $t$, FFT the two-dimensional Fast Fourier Transform (FFT) operator that converts real-space coordinates $(x, y)$ into reciprocal-space wave vectors $(q_x, q_y)$. $\langle\cdot\rangle_{t,q}$ indicates averaging in time and the azimuthal angles of wave vectors. This image structure function corresponds to the theoretically tractable intermediate scattering function of the Brownian particles $f(q, \Delta t)$ by the relation

$$D(q, \Delta t) = A(q)(1 - f(q, \Delta t)) + B(q) \tag{2}$$

where $A(q)$ is the total scattering amplitude depending on the number of particles in view as well as the optical path and associated point spread function, and $B(q)$ a term that captures the effect of imaging noise.

For sufficiently long videos recorded at adequate frame rates under ideal conditions, the amplitude $A(q)$ and noise floor $B(q)$ are revealed by plateaus in $D(q, \Delta t)$ at the largest and smallest time delays, respectively. Without making an *a priori* assumption about the form of $f(q, \Delta t)$, $A(q)$ and $B(q)$ are commonly estimated by fitting these plateaus. For a material with high viscoelastic moduli, it can be prohibitive to wait for tracers to naturally de-correlate and reach the plateau at the largest measured time lag. In addition, slow drifts due to platform instability can also contaminate the shape of the $D(q, \Delta t)$ curve by introducing premature de-correlation as indicated by the orange highlight of the simulated measurement data (larger error bar). Finally, we limited video duration to the same order as the time needed to locate a viable region of interest, typically 20 s per field for mucus measurements; the corresponding acquisition settings are detailed in Section 1.4.

For fluorescence images in which tracer density remains constant and time-independent additive backgrounds are negligible, $A(q) + B(q)$ can instead be calculated from the static image spectrum.[9,10]

$$A(q) + B(q) = 2 \left\langle |\mathrm{FFT}[I(x, y, t)]|^2 \right\rangle_{t,q}. \tag{3}$$

Equation (3) estimates $A(q) + B(q)$ directly, after which $A(q)$ is obtained by subtracting the fitted $B(q)$. Under the assumptions above, $A(q) + B(q)$ equals the fully decorrelated ISF level $D(q, \infty)$, but it is calculated from the image power spectrum and does not require the measured ISF to reach a long-lag plateau. This static-spectrum normalization can improve MSD recovery, but it is not universally unbiased. Its accuracy still depends on the noise estimator, accepted $q$ range, tracer density and intensity, particle size, and imaging scale.[10]

Similar to this approach, we estimated the sum of $A(q)$ and $B(q)$ directly by formally calculating the image structure function based on the difference of a periodically translated frame with itself. While this requires filtering at a specific azimuthal wave number depending on the direction and amount of the translation, we found that fitting to the lower envelope can recover the same estimate but without computing averages of all imaged frames. This leaves the baseline noise floor $B(q)$ as the remaining fitted quantity. Under the independent additive-noise model analyzed by Gu et al., the mean noise term is common across $q$ and lag, although its variance is not.[10] We therefore estimated $B$ from the shortest-lag behavior at high wave number and imposed a common value across the accepted $q$ range only after the consistency search.

Lastly, we can compute the 2D mean squared displacement (MSD) of the tracer particles per wave number $q$ by the relation

$$\langle \Delta r^2 \rangle = -\frac{4}{q^2} \log\left(1 - \frac{D(q, \Delta t) - B(q)}{A(q)}\right). \tag{4}$$

In perfect conditions, the resulting MSD should not be a function of $q$; some algorithms minimize the residual at different $q$ to estimate $A(q), B(q)$.[7] For robustness, we calculated viscoelastic moduli from the median and 10th to 90th quantiles of the obtained MSD over the accepted $q$ range, nominally between 5 and 10 multiplied by objective NA (e.g., $q \in [4, 8]$ for NA $= 0.8$), where the amplitude function $A(q)$ had the expected increasing form.[10] The resulting $q$-wise ranges are descriptive dispersion measures and do not treat wave numbers as independent experimental replicates.

The elastic storage modulus $G'(\omega)$ and viscous loss modulus $G''(\omega)$ are related to the MSD by the generalized Stokes-Einstein relation

$$\begin{aligned}
\alpha(\omega) &= \left.\frac{\mathrm{d}\ln\langle \Delta r^2(\Delta t)\rangle}{\mathrm{d}\ln \Delta t}\right|_{\Delta t = 1/\omega}, \\
|G^*(\omega)| &= \frac{4k_B T}{6\pi r \langle \Delta r^2(1/\omega)\rangle \Gamma[1 + \alpha(\omega)]}, \\
G'(\omega) &= |G^*(\omega)| \cos\left[\frac{\pi\alpha(\omega)}{2}\right], \\
G''(\omega) &= |G^*(\omega)| \sin\left[\frac{\pi\alpha(\omega)}{2}\right].
\end{aligned} \tag{5}$$

where $\omega = 1/\Delta t$ is the frequency, $k_B$ is the Boltzmann's constant, $T$ the measurement temperature, $r$ the hydrodynamic radius of the tracers, and $\Gamma[\cdot]$ the Gamma function. Here $\alpha(\omega)$ is the local logarithmic slope of the MSD evaluated at the lag $\Delta t = 1/\omega$.

The results presented here were obtained using our custom MATLAB implementation of the image-structure-function calculation, with straightforward parallelization and vectorization. In preliminary tests, open-source implementations such as fastDDM accelerated this computationally dominant step by approximately 10–50-fold, depending on the hardware and use of GPU acceleration.[11] Such acceleration could reduce the reported processing time of approximately 20 min per sample to the order of minutes, comparable to the time required for sample preparation.

#### 2.2 Sensitivity of $B(q)$ estimation

For each image sequence, $D_\infty = D(q,\infty) = A(q) + B(q)$ was either measured directly when a long-lag plateau was observed or estimated from Equation (3) or the shifted-image calculation when the plateau was not reached. At a given lag, let $D = D(q,\Delta t)$, $B = B(q)$, $S = D_\infty - B = A(q)$, $x = (D-B)/S$, and $L = -\ln(1-x)$; the two-dimensional MSD is then $M = 4L/q^2$. The accepted range $0 < x < 1$ is the physical condition for a positive, finite, real-valued MSD: $x \geq 1$ makes the logarithm undefined or complex, whereas $x < 0$ gives a negative MSD and $x = 0$ gives zero MSD with divergent relative sensitivity.

We evaluated conditional sensitivity by perturbing $B$ while holding the measured $D$ and selected $D_\infty$ fixed. For comparison, we also perturbed $D_\infty$ separately while holding $D$ and $B$ fixed. To first order,

$$\frac{\delta M}{M} \simeq -\frac{1}{L}\frac{\delta B}{S} - \frac{x}{(1-x)L}\frac{\delta D_\infty}{S}. \quad (6)$$

Thus, an error in $B(q)$ has the greatest effect before appreciable decorrelation, whereas an error in $D(q,\infty)$ has the greatest effect close to full decorrelation. At fixed tracer radius and local MSD slope, the resulting fractional change in $|G^*|$ has approximately the same magnitude and opposite sign because $|G^*| \propto M^{-1}$. For Figure S1, the same $\pm 1\%$ perturbation in $B(q)$ was propagated through the MSD inversion and then through the full generalized Stokes–Einstein conversion, including the perturbation-induced change in the local MSD slope, to obtain $G'(\omega)$, $G''(\omega)$, and $G''/G'$. Although Figure S1 is only a demonstrative example, the amplification of error in $B(q)$ estimation seem reasonably conserved, and should in general not comparable to uncertainties typically seen due to heterogeneity within sample and between replicates for human mucus.

No single method of estimating $B(q)$ and $D(q,\infty)$ is uniformly reliable across all dynamical and imaging regimes.[10] Our four material classes therefore determine how these quantities are obtained. When the ISF physically reached a long-lag plateau, $D(q,\infty)$ was measured directly rather than estimated from Equation (3). When no plateau was resolved in slow or stiff samples, $A(q) + B(q)$ was instead estimated from the static-spectrum or shifted-image calculation and combined with a separately fitted short-lag $B(q)$. This branching avoids fitting a limit that is not resolved within the acquired time window while retaining direct plateau measurements when they are available. Together with the accepted $q$ range and removal of values near $x = 0$ or $x = 1$, it limits amplification of errors in $B(q)$ and $D(q,\infty)$, but does not by itself define a confidence interval.

### 3 Other practical considerations

#### 3.1 Humidification during cell culture

Despite the accessibility provided by DDM, mucus extraction and handling still need further standardization during *in vitro* cell culture maintenance. For our differentiation media tests (Figure 3), we followed published guidance to humidify our cultures every 48 hours and perform apical wash two days before collection to ensure we measure only freshly secreted mucus.[12] We chose to humidify our culture with 50 µL/cm$^2$ of insert area because at lower values, we frequently observed that the apical surface dried out, especially for PneumaCult cultures; this value is, however, not universal. A previous study found that only 3 µL of medium was enough to humidify the cultures.[13] Another study did not specify if a humidifying liquid was used at all, and the mucus was collected only after being loosened 24 hours before collection.[14] These different protocols could be due to either the specific incubator environment or cell donors. Importantly, however, while the collection method will impact the viscoelastic properties of mucus, our results show that relative differences between samples are conserved, as long as the method is not destructive and is standardized within a given experiment.

#### 3.2 In-situ imaging

For in-situ measurements realized via cinnamaldehyde induced ciliostasis, it remains to be verified that such manipulation does not incur long-term damage to the culture, as that would limit the ability to perform longitudinal studies. In our study, only some of the control pHAEC donors (9439) recovered to normal CBF and mucociliary clearance 5 hours after 30 µM cinnamaldehyde treatment for 1 hour through the basal chamber. Tests under diseased conditions, especially those composed of iPSC cells, could not show proper recovery of ciliary activity even at much reduced cinnamaldehyde concentration at 10 µM for less than 20 minutes.

#### 3.3 Stability of mucus in storage

In order to share samples between laboratories and streamline batch analysis, it would be beneficial to store mucus samples for extended periods of time without altering their mechanical properties. Based on published recommendations, we stored mounted

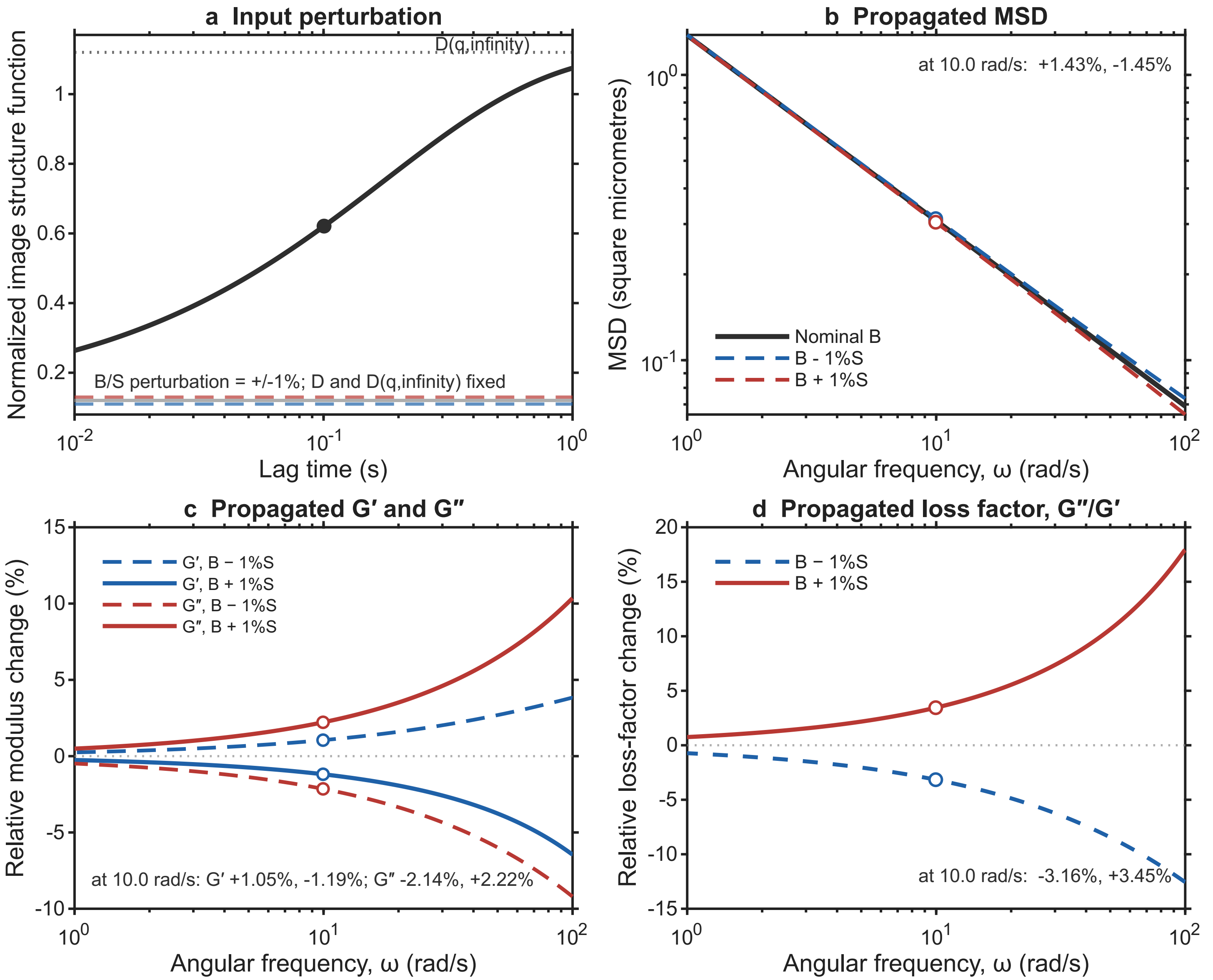


Figure S1: **Propagation of $B(q)$-estimation uncertainties through the DDM rheology conversion. A.** A $\pm1\%$ perturbation in $B(q)$, normalized by the available signal range $S = D(q,\infty) - B(q)$, was applied while holding the measured $D(q,\Delta t)$ and selected $D(q,\infty)$ fixed. **B.** The resulting perturbed MSD curves. **C.** Relative changes in $G'(\omega)$ and $G''(\omega)$ after passing each perturbed MSD curve through the same local-slope and generalized Stokes–Einstein conversion. **D.** Relative change in the loss factor $G''/G'$. Markers and annotations quantify the propagated changes at approximately $10\ \mathrm{s}^{-1}$.

A generic power-law MSD with $\alpha = 0.65$, $q = 3\ \mu\mathrm{m}^{-1}$, and $x = 0.5$ at approximately $10\ \mathrm{s}^{-1}$ was used for this case study.

mucus samples in Parafilm wrapped slides at 4°C.[15] We confirmed that under these conditions, the viscoelastic moduli of extracted *in vitro* mucus remained stable for at least one week, i.e., significantly longer than the typically assumed working time without cryogenic storage for sputum analysis, where contamination is difficult to avoid.[16] See overlapped markers in Figure S2A. Interestingly, mucus from our bronchial airway donor 7783 shows a higher storage modulus than the loss modulus near the typical ciliary beat frequency of about 10 Hz, while the opposite is true for the small airway donor 8938. This indicates that cilia from the bronchial donor are beating against a viscoelastic gel instead of a viscoelastic fluid. As a further proof-of-concept, we show that the fold-change of absolute viscosity for the two donors remained stable between the collection day (Figure S2B, left box plot) and 7 days after storage (Figure S2B, right box plot). This indicates that our measurement and storage protocol can be used to robustly analyze previously stored samples, facilitating batch processing of samples collected at different time points.

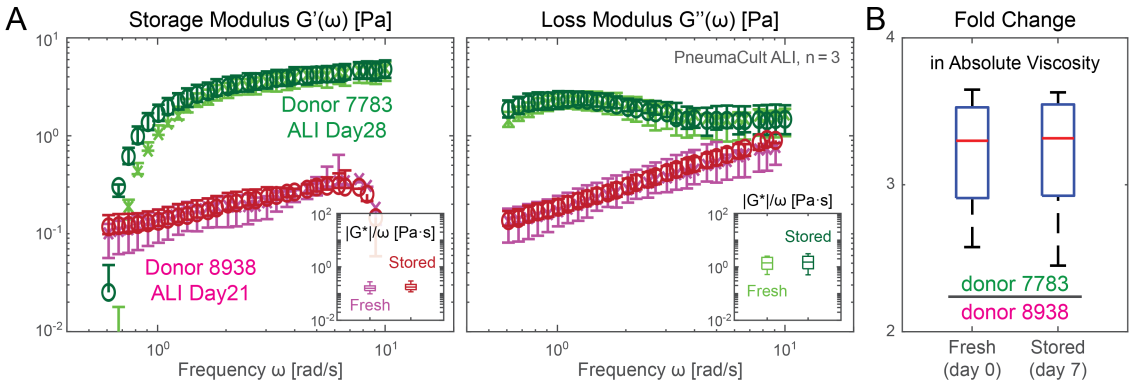


Figure S2: **Rheological stability of sterile *in vitro* mucus collection.** **A.** Storage and loss moduli of mucus extracted from ALI culture maintained in PneumaCult from two separate donors: donor 7783 (green shaded markers) is of bronchial origin, and donor 8938 (magenta shaded markers) is from a small airway source. **B.** Left panels show the fresh and stored absolute viscosity ($|G^*|/\omega$) before (lighter shade) and after 7 days of storage (darker shade). Right panel shows that fold changes in absolute viscosity between the two donors remain stable after 7 days of storage inside the sealed capillary chambers. Measurements performed on pooled $n = 3$ 12-well plate inserts with specified $N = 2$ donors using 3 mm capillary chambers.

#### 3.4 Hydrodynamic radius calibration

The three accepted trials were compared using their own $\omega = 1/\Delta t$ vectors and the common 30–100 [rad s$^{-1}$] window. In Trial #1, Trial #2, and Trial #3 order, the median effective radii were 289.5, 297.6, and 298.2 nm, giving a three-trial mean of 295.1 nm and an observed range of 289.5–298.2 nm. Each trial comprised three raw FOV movies. The wider $q$-wise band in Trial #2 describes dispersion among accepted $q$ modes and was not interpreted as a confidence interval. Figure S3 summarizes the frequency-resolved and post-calibration results.

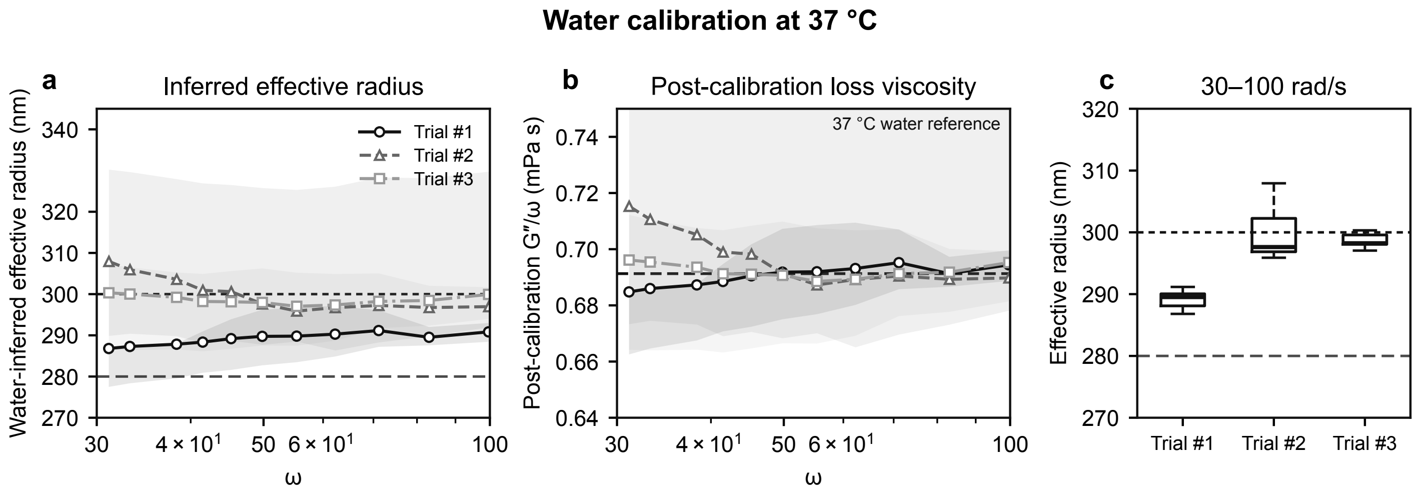


Figure S3: **Water calibration tests at 37 °C.** **A.** Water viscosity inferred effective radius for Trial #1–#3 over 30–100 [rad s$^{-1}$]; shaded bands show the saved $q$-wise 10th–90th ranges. **B.** Post-calibration $G''/\omega$ after applying one trial-wide radius factor. The dashed line is the literature water viscosity. **C.** Boxplots of the frequency-resolved effective radii.

##### 3.5 Validating DDM using synthetic viscoelastic material

Numerous prior studies have demonstrated that DDM microrheology can reproduce correct microrheology results for homogenous standard fluids such as glycerol water mixture and polymer solutions, such as Poly(ethylene oxide) (PEO).[17] To confirm that our high-throughput pipeline is performing on par with expectation, we compared DDM and MPT measurement of 0 to 4% (w/w) PEO solution in Milli-Q water.

51.4 mg of dry PEO powder (900 kDa; Sigma-Aldrich, 189456) were used to make 4% w/w stock solution at room temperature. A total of 1.284 mL of Milli-Q water was gradually added to the powder and mixed until no visible bubbles and chunks were present. The final weight was verified with a digital scale to take account of evaporation during mixing. Dilution series and the stock solution were then stored overnight at 4°C to allow material settling. Before loading 10 $\mu$L samples into capillary chambers, solution in Eppendorf tube was vortexed for homogenization.

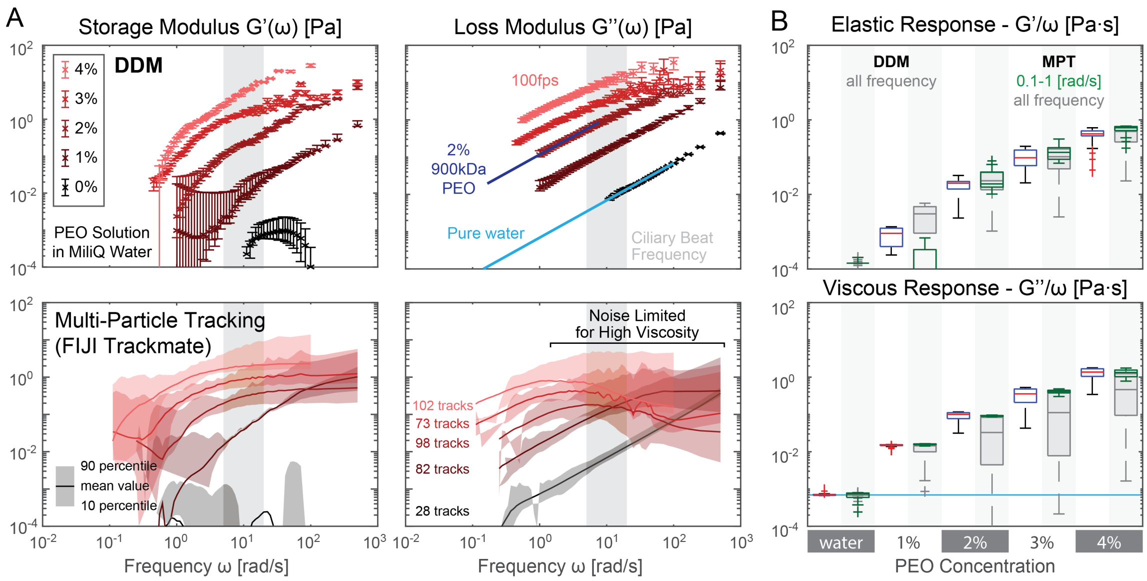


Figure S4: **High-throughput DDM validation with PEO solutions. A.** storage and loss moduli from DDM and particle tracking microrheology. DDM markers show the median and error bars show the 10th–90th $q$-wise range after aggregation across five analyzed ROIs per concentration. These are descriptive ranges, not confidence intervals. The distributions distinguish PEO solutions at different concentrations by weight, especially near typical ciliary beat frequency ranges (light blue). **B.** Comparison of frequency normalized viscoelastic moduli derived from DDM and particle tracking. Particle tracking results closely match that of DDM but only if the noisy frequency range is ignored. DDM box plots are based on all measured frequencies, green MPT box plots are restricted to data measured between 0.1 to 1 Hz, and dark gray for all measured MPT frequencies.

Figure S4 compares the storage and loss moduli of the PEO solution based on our DDM versus MPT measurements. We found an excellent match between the two, especially for normalized loss modulus viscosity, if we restrict the frequency range to 0.1–1 Hz (bottom right panel, green box plots). In Figure S4A, we observe a lower sensitivity bound for storage modulus via DDM below 0.01 Pa. This degradation in performance could be due to edge effects caused by particles moving in and out of view (256$\times$256 px or 166.4$\times$166.4 µm ROI chosen for analysis throughput) for fluids of low elastic response.[18] For similar reasons, only 28 high quality tracks that spanned the full length of the video were found to be analyzed via MPT. Moreover, background artifacts from retarded particle motion near chamber boundaries far away from the focal plane could still contribute some motion for transparent solutions.

#### 4 Biological heterogeneity of human mucus samples

Using COPD donor measurements as an example, Figure S5 shows that variation among ROIs and across technical and biological replicates generally exceeded the $q$-wise dispersion within individual analyses. We reconstructed the COPD storage and loss moduli at $\omega = 1$ and 10 [rad s$^{-1}$] and plotted them against ciliation age. Ciliation age was calculated by subtracting the donor-specific ciliation-onset day from the corresponding ALI sampling day. As shown in Figure 4, ciliation onset occurred on ALI days 35, 33, and 28 for Donors #1, #2, and #3, respectively.

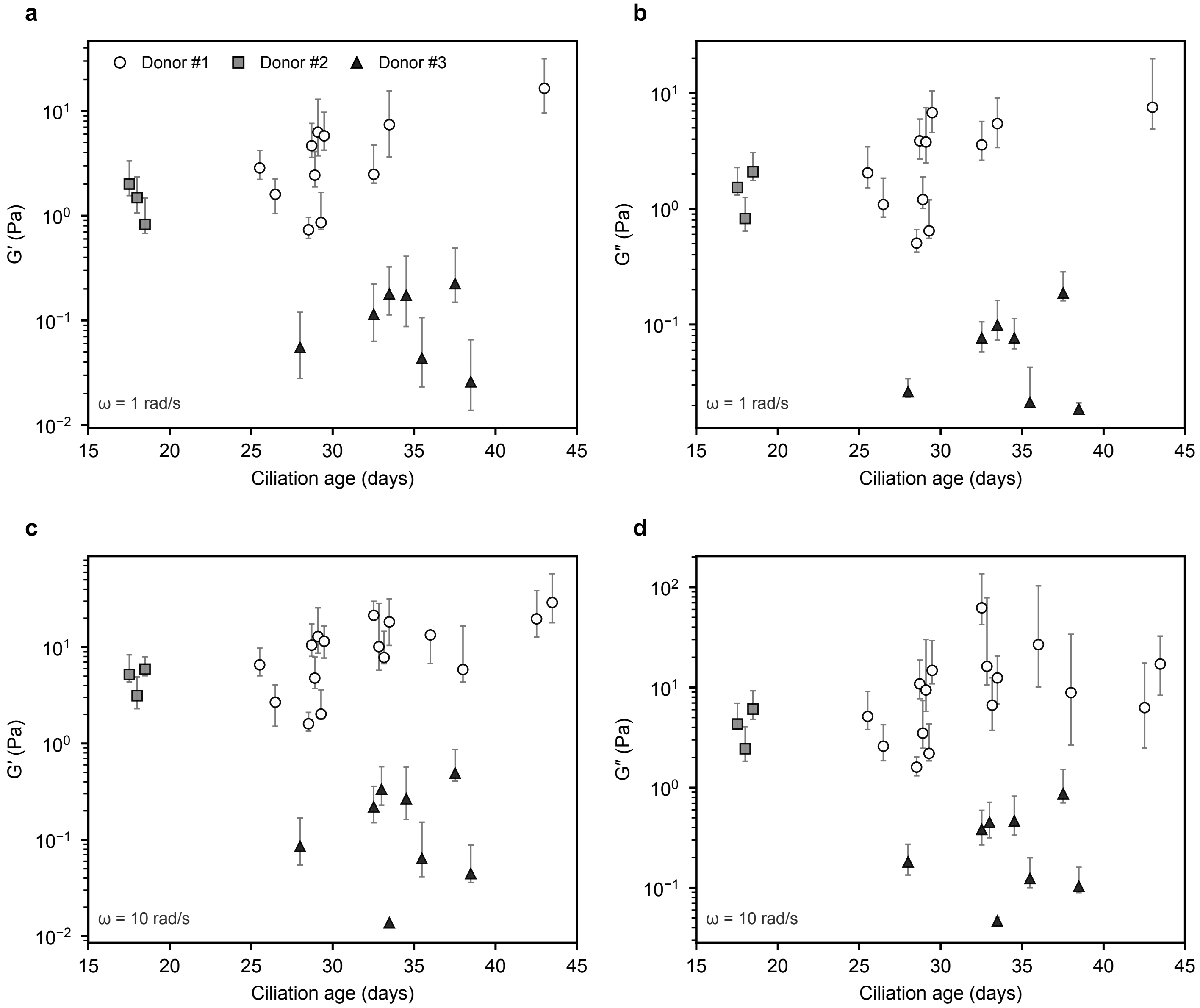


Figure S5: **Observed variation in COPD donor mucus viscoelastic measurements across ciliation ages and frequencies. A,B.** Storage and loss moduli at $\omega = 1$ [rad s$^{-1}$]. **C,D.** Corresponding moduli at $\omega = 10$ [rad s$^{-1}$]. Each marker represents the median from one analyzed ROI or technical replicate, plotted at the corresponding ciliation age. Small horizontal offsets separate observations obtained at the same age. Gray vertical lines indicate the saved 10th–90th percentile range across $q$ values within each analysis. At 1 [rad s$^{-1}$], data were available for 21 observations analyzed using the common reference calibration (Donor #1, $n = 11$; Donor #2, $n = 3$; Donor #3, $n = 7$). At 10 rad,s$^{-1}$, all 27 observations were available (Donor #1, $n = 16$; Donor #2, $n = 3$; Donor #3, $n = 8$). These ranges describe within-analysis $q$-wise dispersion and are not confidence intervals for biological variability.
